# Citizen-Centered, Auditable, and Privacy-Preserving Population Genomics

**DOI:** 10.1101/799999

**Authors:** Dennis Grishin, Jean Louis Raisaro, Juan Ramón Troncoso-Pastoriza, Kamal Obbad, Kevin Quinn, Mickaël Misbach, Jared Gollhardt, Joao Sa, Jacques Fellay, George M. Church, Jean-Pierre Hubaux

**Affiliations:** Department of Genetics, Harvard Medical School, Boston, USA; Precision Medicine Unit, Lausanne University Hospital, Lausanne, Switzerland; School of Computer and Communication Sciences, École Polytechnique Fédérale de Lausanne, Lausanne, Switzerland; Nebula Genomics Inc., San Francisco, USA

## Abstract

The growing number of health-data breaches, the use of genomic databases for law enforcement purposes and the lack of transparency of personal-genomics companies are raising unprecedented privacy concerns. To enable a secure exploration of genomic datasets with controlled and transparent data access, we propose a novel approach that combines cryptographic privacy-preserving technologies, such as homomorphic encryption and secure multi-party computation, with the auditability of blockchains. This approach provides strong security guarantees against realistic threat models by empowering individual citizens to decide who can query and access their genomic data and by ensuring end-to-end data confidentiality. Our open-source implementation supports queries on the encrypted genomic data of hundreds of thousands of individuals, with minimal overhead. Our work opens a path towards multi-functional, privacy-preserving genomic-data analysis.

**One Sentence Summary:** A citizen-centered open-source response to the privacy concerns that hinder population genomics, based on modern cryptography.

---

The decreasing costs of DNA analyses and the promise of the value of large genomic datasets have ushered us into the era of population-scale genomics. This development is driven by companies, as well as governments. The direct-to-consumer (DTC) personal-genomics market has grown dramatically over the past few years (*1*) and multiple countries around the globe are pursuing large-scale population-genomics initiatives. The growing genomic datasets promise to improve preventive medicine and to support the development of more targeted therapies.

This potential can be realized only if genomic data is made easily and widely accessible to researchers. Unresolved privacy issues, however, make data sharing on a large scale extremely difficult and time-consuming. Historically, health-data privacy has always been addressed by data “de-identification”, specifically by removing the fields that reveal an individual’s identity. However, personal genomic data cannot be effectively de-identified, because even a small subset of it is sufficient to identify an individual or a relative, as illustrated notably by the successful usage of DNA in forensics (*2, 3*). People are particularly sensitive and concerned about genomic-data privacy, mainly because of the real or perceived risk of genetic discrimination in areas such as insurance, employment, and education. Although it is not possible to determine whether such concerns are fully justified, it has been established that they deter individuals from participating in population-genomics studies and from using direct-to-consumer genetic testing services. We surveyed 442 individuals interested in genetic testing and found that data privacy (Supplementary Materials Section 2.6) is one of the top two disincentives, together with cost (Figure 1A). In particular, individuals want to have control over the use of their genomic data and to be able to share access to their data with researchers, without risking misuse; these are guarantees that population-genomics initiatives and personal-genomics companies are not able to provide today (Figure 1B).

**Figure 1.**
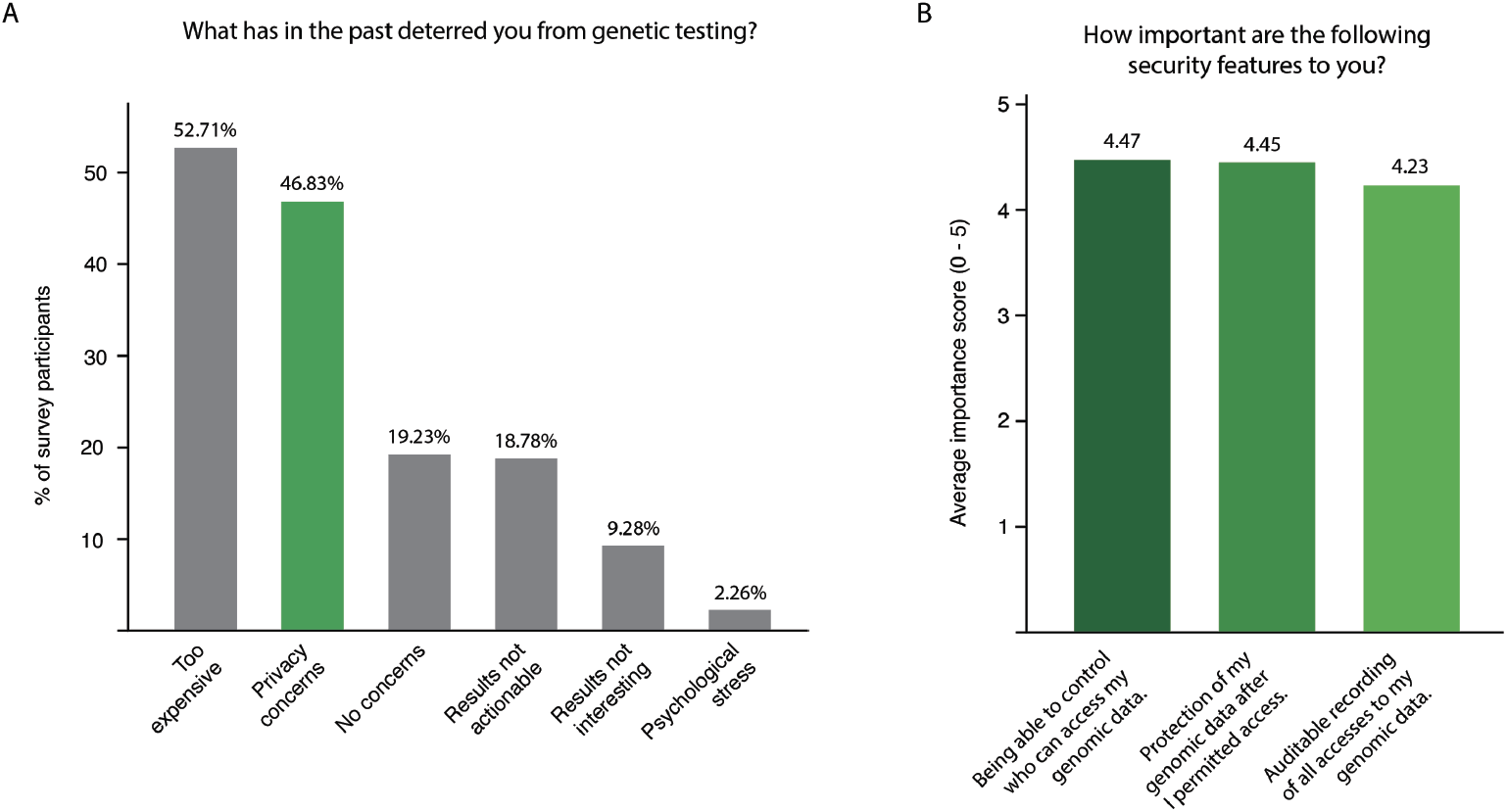
Results of a survey of 442 individuals interested in genetic testing and data sharing. (A) Cost and privacy concerns are prevalent. (B) Survey participants want to control access to their data, to be able to share it securely and the process to be transparent and auditable.

In the last few years, researchers from the information security community have proposed solutions that address these requirements. Some of them focus on providing secure storage for genomic data, whereas others propose methods on how to securely carry out specific computations (*4, 5*). None of these solutions, however, has so far been adopted in practice, mainly because of their limited scope and applicability to real operational use cases. Furthermore, the issue of trust goes beyond the provision of secure storage and processing, as it is also intimately related to the question of transparency and individual control of data sharing and use. This particular need is currently not met by existing policies and genomic-data-sharing platforms. For example, the NIH Genomic Data Sharing (GDS) Policy (*6*) describes a data-sharing model that relies on broad consent policies and institutional governance over datasets. This broad consent policy is adopted by existing genomic-data-sharing platforms, such as the GA4GH Beacon Network, that facilitate genomic-data sharing between institutions, but do not give dynamic, granular control to individuals who provided the genomic data (*7*). However, the NIH and other policymakers recognize that “changing technology can enable more dynamic consent processes that improve tracking and oversight and more closely reflect participant preferences” (*8*).

The solution we propose here is guided by the results of our survey and provides a concrete implementation of some of the ideas for privacy-preserving personal genomic recently discussed in a position paper that involved some of the authors of this work (*9*). By combining several privacy-preserving technologies, including homomorphic encryption, encryption-based access control by data owners, and blockchain technology, our solution enables citizens to share their genomic and clinical data in an encrypted form, through a secure platform that ensures that their data is discoverable by researchers while it remains always encrypted and protected against unauthorized access. The platform also ensures that authorized researchers can access and use decrypted genetic data for further analyses in a data-owner-controlled and auditable way.

More specifically, our platform is the first to simultaneously ensure (i) distribution of trust, (ii) end-to-end protection of genomic data, (iii) secure data release, and (iv) citizen-defined and auditable data control, as explained below (Figure 2):

**Figure 2.**
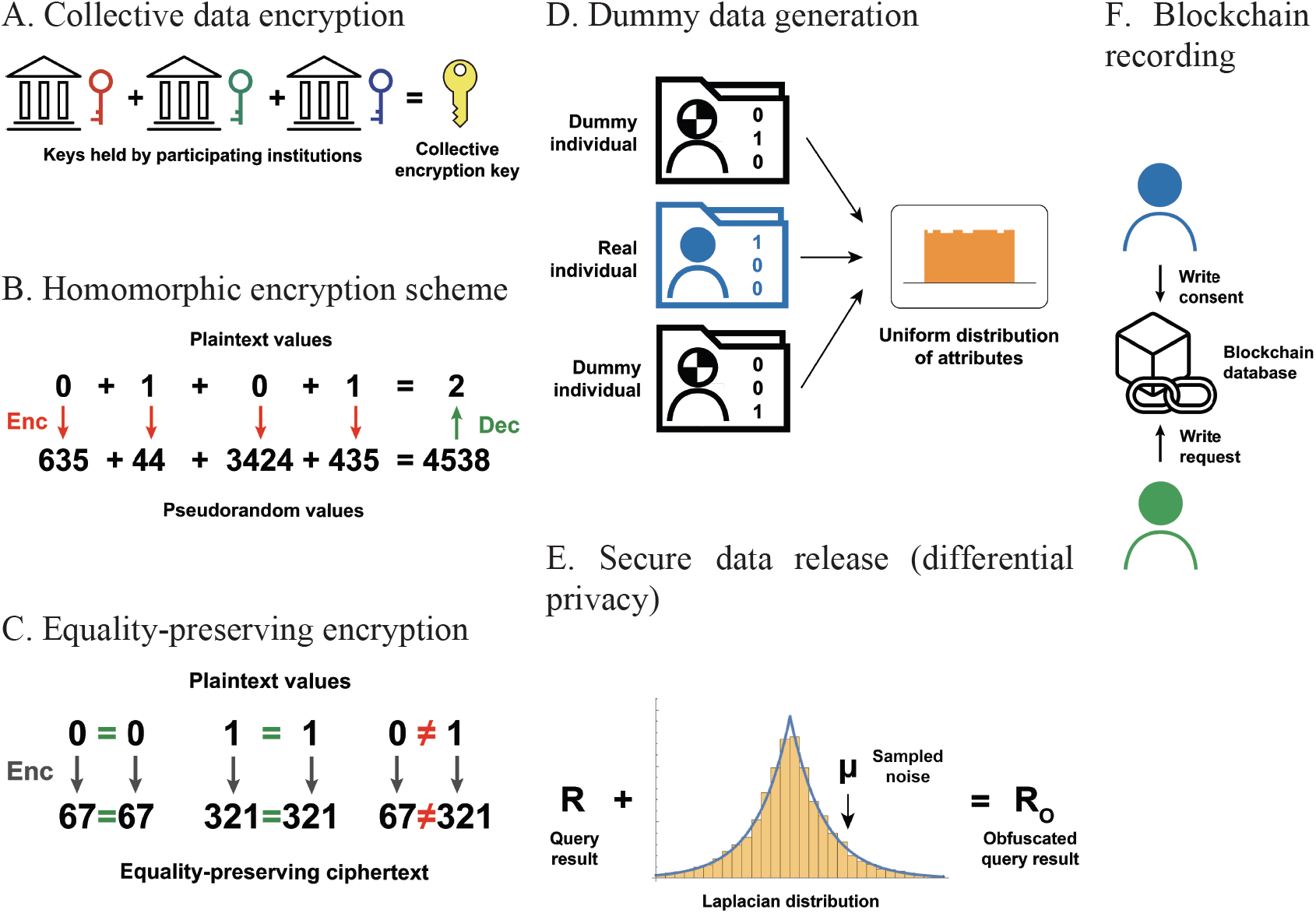
Security and privacy techniques used by the proposed platform: (A) Data is collectively encrypted using multiple keys that are held by independent parties. (B) Attributes such as genotypes are encrypted using an additively homomorphic scheme that enables computing on encrypted data. (C) Equality-preserving encryption scheme enables equality-matching queries. (D) Dummy individuals are added to the system in order to even out the distribution of various attributes and prevent frequency attacks. (E) Noise is added to query results to prevent identification of individuals, based on their combination of attributes. (F) Records of all activities, in particular, data-sharing consents and data-access requests, are immutably stored on a blockchain in order to ensure auditability.

## Distribution of trust

is achieved when the protection of the data on the platform is guaranteed by multiple parties and not by a single one. This is ensured in the proposed platform by encrypting individual’s data with a collective key that is jointly generated by a set of computing nodes operated by independent, but collaborative, parties (e.g., different organizations) (Figure 2A). The use of such encryption key guarantees that citizens/researchers do not need to rely on a single central authority for the protection of their data/queries, because an attacker would need to compromise all the computing nodes in order to obtain the decryption key. Intuitively, the more computing nodes that participate in the generation of the collective key, the more secure the data is. We assume that these nodes can be instantiated by highly available service providers and/or clinical institutions. But if this is not the case, it is also possible to reduce the number of nodes needed to obtain the decryption key to a fraction (threshold) of the total, in order to avoid system disruption if a minority of the nodes goes offline, hence achieving a balance between security and availability.

## End-to-end protection

of data (i.e., during storage, transfer, and computation) is achieved by the combined use of homomorphic encryption and equality-preserving encryption. Homomorphic encryption enables computations on encrypted data without having to decrypt it. For example, in a hospital dataset, patients who are affected by a determined rare disease are labeled by a 1 and those who are not affected are labeled by a 0. If these numbers are encrypted with homomorphic encryption, they will look like indistinguishable pseudorandom numbers, and it will be unfeasible to tell patients apart without the decryption key. However, if the encrypted numbers are summed together, they will produce a result that, when decrypted, corresponds to the number of patients affected by the rare disease (Figure 2B). Whereas, equality-preserving encryption guarantees that numbers encrypted with the same key always generate the same ciphertext (Figure 2C). This type of encryption is necessary to enable equality-matching operations on encrypted attributes, without revealing them; this cannot be efficiently performed with homomorphic encryption. However, if it is not used in the right way, equality-preserving encryption is vulnerable to frequency attacks because of its equality-preserving property. These attacks aim at breaking the encryption by reconstructing the distribution of the encrypted data and mapping it to a known unencrypted distribution. In the aforementioned rare-disease example, where the presence or absence of the disease in each patient is labeled with a 1 or a 0, even if the labels are encrypted, it can be inferred that patients with the encrypted label that appears less often are the ones with the rare disease. To mitigate this type of attack, equality-preserving encryption must be combined with a technique that generates additional dummy data (or patients in this case) such that the frequency differences of the encrypted labels are evened out, thus resulting in a uniform distribution (Figure 2D). More details about homomorphic encryption and equality-preserving encryption are provided in the Supplementary Materials Section 1.1.

## Secure data release

Even when the data is protected end-to-end during the computation, the released aggregate results can also be used by an attacker to try to re-identify individuals whose data was part of the input dataset, through membership or attribute inference attacks (*10*). In the rare-disease hospital-dataset example, an attacker could easily infer with high confidence that a target individual is part of the dataset if he holds some background information about the target, such as demographic data or some clinical attributes that further restrict the set of matching rare-disease patients to a very low count. There are several frameworks (*11, 12*) for quantifying and preventing undesired inferences on the released data (see Supplementary Material Section 1.3), among which the most widespread is differential privacy.^9^ A data-release mechanism, such as a data-discovery query, is differentially private if it is not possible for an attacker to use the query result to infer whether the data of any given individual was used by the query (Figure 2E). In other words, differential privacy guarantees that it is not possible to re-identify any individual from the query result. Differential privacy is ensured in the proposed platform by obfuscating query results with noise sampled from a given probability distribution. Differential privacy also introduces the notion of a privacy “budget” that can be used to limit the number of queries the same researcher can run on the platform thus preventing attackers who would try to maliciously use the platform in order to reconstruct sensitive attributes of individuals.

## Citizen-defined and auditable data-control

is ensured by logging every activity performed in the platform (e.g., data upload, query execution, data-access request) on a permissioned blockchain and by using smart contracts (self-executing protocols that can change the blockchain global state) to automatically execute data-access policies dynamically defined by the data owners. A blockchain is an immutable, collectively maintained database (Figure 2F) to which new entries can be added only if they have been verified and approved by a majority of the computing nodes that maintain the blockchain. Entries are bundled into timestamped blocks and each block references its preceding block, which creates a sequential ordering that prevents malicious deletion or reordering of data stored on the chain. This enables tamper-proof, auditable record-keeping. In addition to data, entries can also contain smart contracts. Note that the permissioned (or private) blockchain requires a thorough definition and rigorous management of the participating computing nodes, thus it is fundamentally different from the more widespread permissionless (or public) blockchains used for cryptocurrencies such as Bitcoin.

Our proposed platform provides these privacy and security guarantees and enables data-discovery analyses and, eventually, access to individual-level genetic data through a secure protocol that synergistically combines all the above-mentioned techniques. This protocol consists of three phases that involve all the parties depicted in Figure 3.

**Figure 3.**
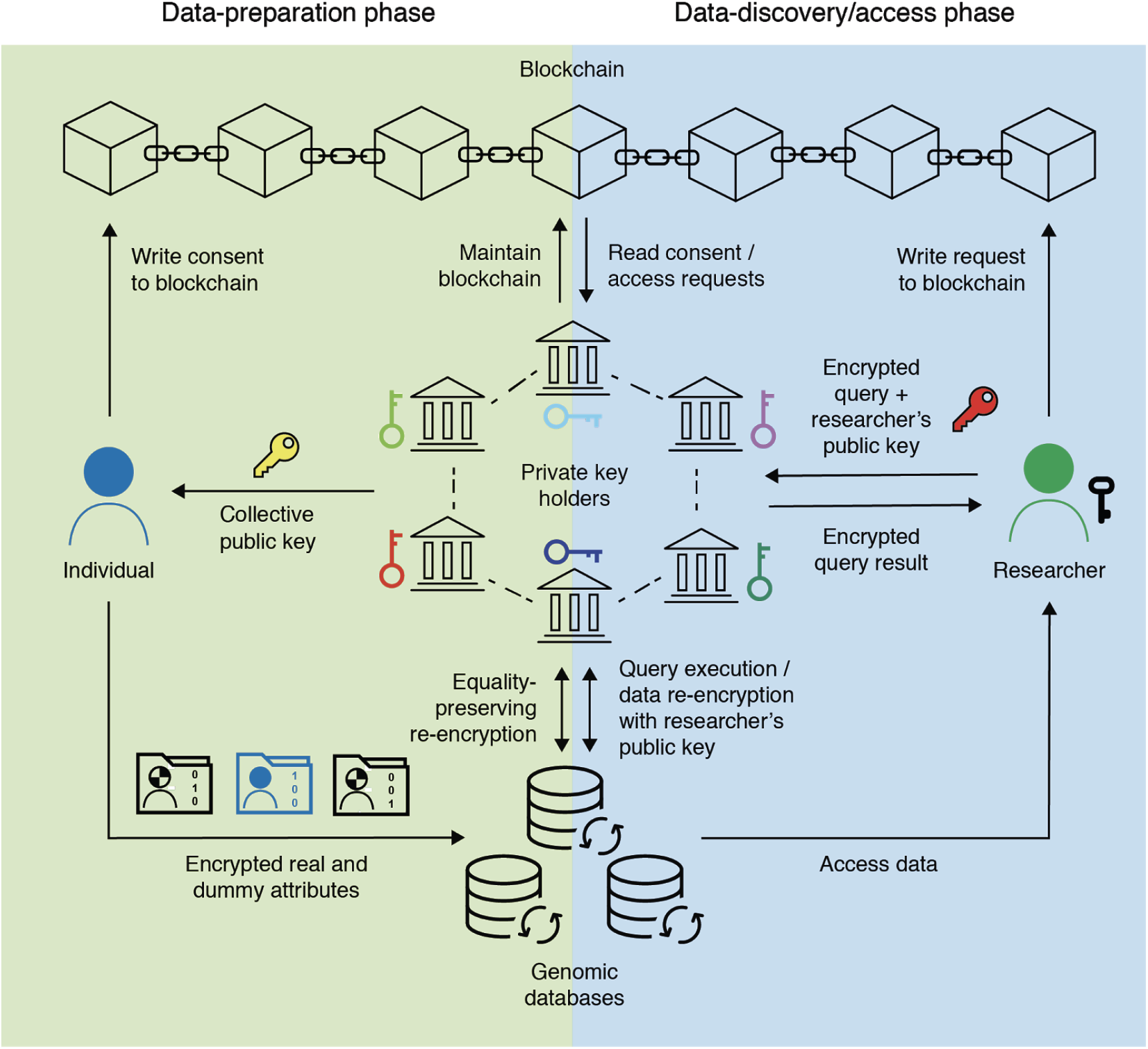
Platform overview with the three phases of the protocol: *data-preparation phase, data discovery phase*, and *data access phase*.

In the *data-preparation phase*, a set of independent, yet collaborative, computing nodes generate one public-private cryptographic key pair each and combine their public keys into a single collective public encryption key for an additively homomorphic encryption system. When an individual wants to upload her data, a set of dummy individuals are generated with specific variants and clinical attributes, chosen according to an estimated population distribution from public data and data already uploaded to the platform, such that the overall distribution of variables within the platform is uniform. Then, each genetic variant and each clinical attribute of the real individual and the dummies is encrypted with the collective public encryption key for the additively homomorphic encryption system. To prevent dummy individuals from affecting aggregate results obtainable during the data-discovery phase and for them to still be indistinguishable from the real individual, a binary flag is appended to each individual (real and dummy) and is also encrypted under the collective public key. The binary flag is set to “0” for dummy individuals and to “1” for the real one. In addition to the individual encryption of various searchable entries, raw genetic and clinical files (e.g., BAM -Binary Alignment Map-files, VCF -Variant Call Format-files, or PDFs of medical records) are encrypted with a self-generated symmetric key that is then encrypted with the collective public key. The data provider also defines a data-access policy that is uploaded and stored together with the data on the platform. Data uploads generate transactions on the platform blockchain that serve as records of data uploads and that immutably associate the data provider’s identity with each file identifier. Finally, in order to enable equality-matching queries, computing nodes run a distributed re-encryption protocol on the uploaded data to switch the encryption of the genetic variants and clinical attributes from the original homomorphic form to an equality-preserving form. The presence of dummy individuals protects from frequency attacks the data encrypted in such form. Further details about the *data-preparation phase* and the generation about the dummy individuals are explained in Supplementary Materials Section 2.3, and Figures S2, S3, and S4.

In the *data-discovery phase*, the researcher can run combinatorial Boolean queries, made of clinical and genetic inclusion/exclusion criteria, on the encrypted data in order to obtain aggregate statistics or the list of identifiers for the individuals matching the search criteria. To this end, the queries are processed in a similar way to uploaded data: The researcher encrypts the query items with the collective public key, and the computing nodes re-encrypt them from a homomorphic to an equality-preserving format and log them on the blockchain for traceability. When queries are run on the encrypted database, the homomorphically encrypted flags of the real and dummy individuals whose attributes match the equality-preserving-encrypted query items are retrieved. If the query requests the number of individuals matching the search criteria, the flags are homomorphically summed (the contribution of dummy individuals to the encrypted sum is null), differentially private noise is added, and the encrypted result is returned. If the query requests the identifiers of the individuals who match the query, the latter are masked with the encrypted flags and returned in encrypted form. The results are re-encrypted by the computing nodes with the public key provided by the researcher and sent back to her. The researcher can now decrypt the results with her private key. After decrypting the identifiers, the real individuals are separated from the null values (dummies). This process is transparent to the researcher, and it happens in the front-end module of the platform. Further details about the *data-discovery phase* are explained in Supplementary Materials, notably Figure S5.

In the *data-access phase*, researchers who need to conduct data analysis beyond data-discovery queries, e.g., to conduct genome-wide association studies (GWAS) in order to discover novel variants, can access cleartext individual-level data in a transparent, auditable fashion that ensures accountability. In particular, the researcher can request access to raw files for individuals identified during the *data-discovery phase*. The request is logged on the blockchain, as well as matched against the access policy for the requested files. There are three possible outcomes for such a request: either file access is granted, or it is rejected, or the individual is requested to review and approve the specific request. If file access is granted, the symmetric key that was used to encrypt the file during the data-preparation phase is re-encrypted with the researcher’s public key and sent back to her. The researcher can decrypt the symmetric key and use it to decrypt the file. Further details about the *data-discovery phase* are explained in Supplementary Materials, notably Figure S6.

To demonstrate the practicality of our privacy-preserving approach for population genomics, we implemented an operational prototype of the platform and benchmarked it on a simulated dataset of 150,000 individual records with a total of 28 billion genetic variants (more details on the dataset are provided in the Supplementary Materials Section 3.1). The source code of our prototype is publicly available on GitHub.^1^ The performance results in Figure 4 show that a researcher can discover individuals from such dataset in a few minutes, with queries involving genetic and clinical attributes over the encrypted space; furthermore, the query response time grows linearly with the number of individuals and query items (clinical and genetic criteria used to build the query), whereas the overhead introduced by the encryption with respect to the clear text is negligible (a few seconds), linear in the size of the matching set, and virtually constant for the database size or number of query items. Even large queries that consist of hundreds of items are executed in just a few minutes with an average overhead of only 1% to 3%, compared to querying plain-text data. Benchmark results for the preparation phase are illustrated in Supplementary Figures S7 and S8, and Figure S9 and show the breakdown for the query-response time where the database processing-time (unaffected by the encryption) is the largest component. Importantly, although the preparation phase is relatively time-consuming, it must be executed only once for every new data provider who uploads data. Finally, the efficiency of the *data-access phase* is determined by the blockchain performance, as access to each file requires at least one transaction, access being regulated by individual consent. The blockchain implemented as part of our system achieves a throughput of thousands of transactions per second (Supplementary Figure S10).

**Figure 4.**
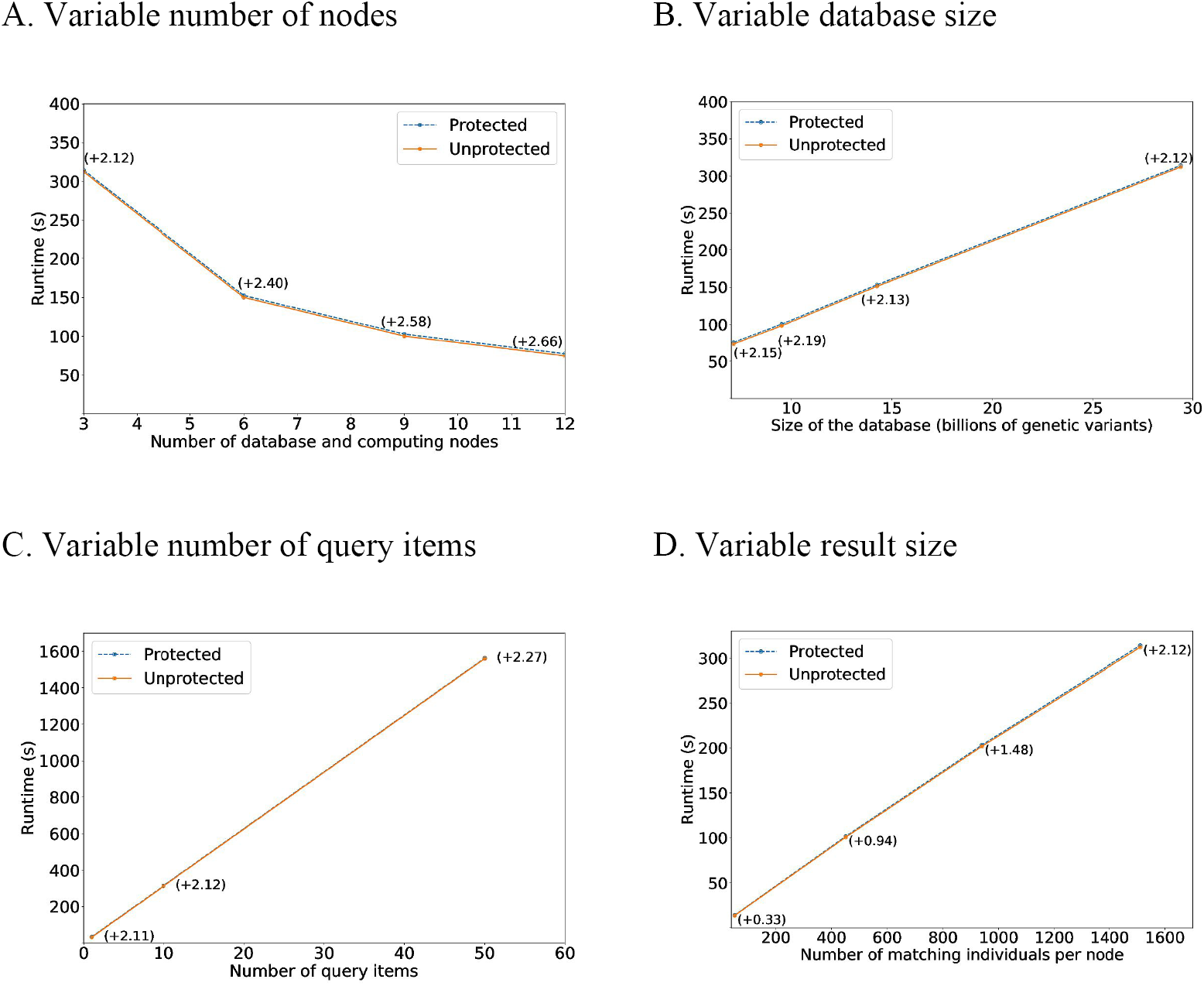
Performance of the data discovery, with a total of 150,000 individuals, each one with a number of observed variants that ranges from 15,000 to 200,000, when (A) varying the number of database and computing nodes for a fixed database size of 28 billion genetic variants, and a query with 10 items producing a resulting set of 1511 individuals per node (worst-case scenario on the database response time; a constant total size for the resulting set would imply a faster decrease of the response time), (B) varying the size of the database for 3 database and computing nodes, a query with 10 items and a resulting set of 1511 individuals per node, (C) varying the number of query items for 3 nodes, a database of 28 billion observations and a query returning 1511 individuals per node, (D) varying the size of the returned matching set for 3 nodes, a database of 28 billion genetic variants and a query with 10 items. The difference between the *protected runtime* (meaning that operations are carried out on encrypted data) and *unprotected runtime* (meaning that operations are carried out on cleartext data) is shown in parenthesis for each data point, in seconds. This difference is the data-protection overhead.

The efficient, privacy-preserving, and auditable data-discovery functionality provided by the proposed platform can enable a number of common use cases for genomic data on millions of individuals with response times in the order of a few minutes. For example, it enables drug-target validation whereby researchers test whether a genetic variant associated with a drug target is more common in a case group than in a control group for different populations spread out over the entire globe. The platform can also be used to determine frequencies of genetic variants of interest in the population before recruiting for clinical trials. This can help avoid failures in recruiting due to too narrow inclusion criteria. The querying of clinical data is also useful in the absence of genomic data. For example, it can enable the identification of a cohort with a phenotype of interest that can be subsequently sequenced at the expense of interested researchers. In this scenario, enabling individuals to receive sponsored genome sequencing without having to reveal any personal information in advance would help to overcome the price barrier that is still hindering many people from benefiting from personal genomics (Figure 1A).

In summary, our work advances the field in three ways. First, whereas most of the previous studies focused on the application of individual cryptographic techniques, this is the first solution that integrates multiple complementary technologies that, when combined, enable the controllable, transparent and secure sharing of genomic data. Second, whereas previously proposed implementations focused on institutional genomic data sharing, our solution is citizen centric. The design of the proposed platform was guided by the users’ demand for personal genomics services and it is built to put them in control of their personal genomic data. Third, our implementation, which is open-source and available on GitHub, achieves practical performance at an unprecedented scale with strong privacy and security guarantees. We demonstrate that the genomic data of hundreds of thousands of individuals can be securely explored by researchers with minimal computational overhead due to encryption.

Furthermore, our approach is general and portable. It can be easily integrated with other secure computation protocols that have been developed over the past few years for more specific and (mostly) centralized bioinformatic analyses, thus making them more secure (by decentralizing trust across multiple nodes) and more citizen-centric (by enabling individuals to have control over the use of their data). For example, the functionalities of the proposed platform can be complemented by also enabling data providers to securely gain insight from their personal genomic data. This can be best accomplished by providing polygenic risk scores or by comparing one’s personal genome with as many other genomes as possible, for example to identify genetic variants that are associated with a certain trait or disease. Jagadeesh *et al.(5)* describe a privacy-preserving protocol for genomic comparison in a manner that minimizes the information exposed about all the participants’ genetic information. Similarly, McLaren *et al.(13)* show how to compute polygenic risk scores on homomorphically encrypted genomes. Our platform can also be expanded by enabling researchers to perform further analytical tasks on selected cohorts, without having to request full data access from the individuals. For example, Cho *et al.(4)* demonstrate that a secure multi-party computation protocol can be optimized for GWAS and enable practical performance at a large scale. If integrated with our system, a protocol such as this would further minimize privacy risks and encourage more people to share their personal genomic data with researchers.

Genomic-data privacy has been a very active area of research. However, because of misaligned interests, so far there has been almost no real-world adoption of the developed privacy-preserving technologies. Although individuals welcome the utilization of cryptographic techniques, as our survey has shown, these techniques can also be perceived as a hindrance by researchers. We have shown, however, that the performance and functionality of a privacy-focused platform can be on par with those of unprotected systems, therefore debunking this perception and yielding a beneficial solution both for individuals, whose privacy and control over data is fully respected and auditable, and for researchers, who can have controlled access to larger datasets that would otherwise not be discoverable. We believe that it is time to test this hypothesis by integrating privacy-preserving protocols that have been developed over the past years into a fully functional system. With the approach and implementation we describe here we have taken the first step in this direction.

## Acknowledgments

We are grateful to the Bitfury Exonum engineering team for running part of the performance measurements described in the paper. We also thank Claire Redin from Lausanne University Hospital and John Aach from Harvard Medical School for their useful feedback and constructive comments on the manuscript.

## Funding

This work was supported in part by the grant #2017-201 (DPPH) of the Swiss strategic focus area Personalized Health and Related Technologies (PHRT), and by the grant #2018-522 (MedCo) of the PHRT and the Swiss Personalized Health Network (SPHN). The work was also supported by Nebula Genomics, Inc.

## Author contributions

DG, JLR and JTP have developed the methods. DG, JLR, JTP and JPH wrote the manuscript. DG, JLR, JTP, KO, JG, MM, KQ and JS implemented the algorithms described in the methods and performed the benchmark experiments. All the authors discussed and reviewed the manuscript at all stages. JF, GMC and JPH supervised all the work.

## Competing interests

DG, KO, KQ, JG and GMC are affiliated with Nebula Genomics. JLR, JTP, MM, JS, JF and JPH declare no competing interests.

## Data and materials availability

All data, code, and materials used in the analysis are available on GitHub: https://github.com/ldsec/ccgd-platform

## Supplementary Materials

We begin the supplementary material by introducing a high-level description of the privacy and security concepts used in the paper. We continue by describing in details the material and methods that support our system for secure, privacy-preserving and auditable data sharing. Finally, we report and describe additional performance results and the security analysis of the proposed platform that, due to space constraints, are not included in the main manuscript.

### 1 Privacy and Security Background

We first introduce the main cryptographic concepts used in our platform, with a special focus on privacy and security. In particular, we further develop: (a) secure computing techniques that enable data processing while avoiding undesired leakages to the computing party or parties; (b) data integrity and trust decentralization techniques, that enable immutable storage of relevant logging and transaction information guaranteeing tamper-proof integrity with no single points of failure, and (c) secure data release techniques, that enable sharing cleartext data (such as the results of a secure process) while mitigating malicious attacks on those data (e.g., membership inference or reconstruction attacks).

#### 1.1 Secure computing

Confidentiality and privacy in collaborative processes are traditionally handled by the use of cryptographic privacy-enhancing techniques, especially homomorphic encryption and secure multi-party computation. In particular, our system requires database searches, which are more efficiently handled by equality-preserving encryption. We now give a high-level overview of these three technologies.

##### 1.1.1 Homomorphic Encryption

Homomorphic Encryption (HE) enables computing on encrypted data without having to decrypt it first, consequently being an excellent enabler for secure processing in untrustworthy environments. By virtue of a mathematical homomorphism between the plaintext space and the ciphertext space, a server without access to the secret decryption key can obtain the result of group (addition or multiplication) or ring (addition and multiplication) operations on the plain-text by running their homomorphic counterparts on the ciphertext. Depending on the available homomorphism, HE systems can be either additive, multiplicative or algebraic (both additive and multiplicative). The latter are usually called somewhat homomorphic cryptosystems (SHE) when they enable a limited number of subsequent products, or fully homomorphic cryptosystems (FHE) when they enable an unlimited number of chained homomorphic operations. In order to be considered secure, any homomorphic cryptosystem must feature *semantic security*, which means that it is not possible for somebody without access to the secret decryption key to distinguish between encryptions of the same or different plaintexts. This requires the cryptosystem to be *probabilistic*, i.e., each encryption is randomized so that two encryptions of the same plaintext result in different ciphertexts with very high probability.

Most of the current homomorphic privacy-preserving systems are based on efficient somewhat homomorphic cryptosystems (SHE) relying on the hardness of Ring Learning with Errors (RLWE) lattice problem, such as FV (*14*) and BGV (*15*). These cryptosystems can handle a limited number of arithmetic functions (additions and multiplications) on the ciphertexts without decrypting them, so their versatility is restricted, and they have to be paired with other techniques for them to support functional systems. Recent advances in fully homomorphic encryption (FHE) (*16*), which can handle an arbitrary number of encrypted operations, are promising, but their computational overhead is still far from practical; meanwhile, SHE is now being extensively optimized for achieving better scalability (*17*), and it is also starting to be applied to practical health scenarios, including the support of some basic machine learning primitives, including linear and logistic regression functionalities (*18–24*).

###### Elliptic Curve ElGamal

Our system requires an additively and probabilistic homomorphic cryptosystem to enable counts and histogram generation with encrypted data; for this purpose, we could have chosen either a lattice-based SHE, or an additively homomorphic cryptosystem. In order to (optionally) enable verifiability of the performed operations, we chose the latter, and more specifically, Elliptic Curve ElGamal (*25*), which can support an efficient use of zero-knowledge proofs for correctness, contrarily to lattice-based SHE. Zero-knowledge proofs (*26*) are cryptographic constructions by which a prover can convince a verifier that a statement is true without revealing any further information.

It must be noted, though, that our system functionality is not bound to this choice and can be achieved with other cryptosystems. The encryption of a message *m* ∈ ℤ_*p*_ (the set of integers between 0 and a big prime *p*), is a pair of points in the elliptic curve: *E*_*K*_(*m*) = (*rG, mG*+*rK*), where *r* is a uniformly-random nonce in ℤ_*p*_ (therefore the probabilistic nature of the encryption), *G* is a base point on an elliptic curve *γ*, and *K* is a public key.

The additive homomorphic property states that *E*(*αm*_1_ + *βm*_2_) = *αE*(*m*_1_) + *βE*(*m*_2_) for any messages *m*_1_ and *m*_2_ and for any scalars *α* and *β*. In order to decrypt a ciphertext (*rG, mG* + *rK*), the holder of the corresponding private key *k* (such that *K* = *kG*) multiplies the first point *rG* and *k* yielding *k*(*rG*) = *rK*, and subtracts this point from the second term of the ciphertext *mG* + *rK*. The result *mG* is then mapped back to *m*, e.g., by using a hash table. We rely on fixed-point representation to encrypt floating values.

As described in the main text, we enable trust distribution by combining the public keys of several computing servers and building a collective public key. Decryption is then enabled as an interactive protocol, where each server sequentially performs a partial decryption with its own secret key (which is still protected by the security properties of EC ElGamal, by virtue of the discrete logarithm problem).

##### 1.1.2 Secure Multi-Party Computation

Secure multi-party computation (SMC) is an area of cryptography that aims at enabling several parties to evaluate a function on private data coming from distinct data sources without aggregating or sharing the input data (*27*). At the end of the protocol, the parties learn nothing more but the value of the function. SMC protocols are primarily based on either secret sharing (splitting the secret values in randomly generated *shares* that are distributed among all parties) or garbled circuits (hashing the truth tables of the to-be-computed circuit gates and executing them by obliviously transferring the inputs) (*28*). Recent progress on *oblivious transfer* made a lot of these protocols practical and even scalable (*29*), thus allowing for privacy-preserving machine learning computations. Various privacy-preserving supervised machine learning algorithms have been proposed and analyzed in the SMC setting (*30*). There is an increasing interest in both training and prediction algorithms for machine learning under SMC (*31, 32*).

##### 1.1.3 Equality-Preserving Encryption

The probabilistic nature of homomorphic cryptosystems, where the encryption function is a one-to-many function (it takes one same input to several different possible output values), prevents direct equality comparisons between ciphertexts. There are two options to enable this functionality: either (a) define the comparison function as a polynomial that can be homomorphically executed with the ring operations enabled by the cryptosystem; this solution is computationally complex, especially when the plaintext coefficients are not represented with a binary decomposition. Moreover, the result of the comparison would still be encrypted, requiring decryption or further homomorphic processing to be effectively used. (b) The other option is to use instead a non-probabilistic cipher in which the encryption of the same plaintext will always yield the same ciphertext; i.e., the encryption function is a one-to-one function (deterministic). This enables very efficient comparisons by just comparing the values of the ciphertexts.

In the case of EC ElGamal encryptions *E*_*K*_(*m*) = (*rG, mG* + *rK*), a deterministic version can be achieved by setting a fixed (secret) value for *r*, hence using only the second point of the encryption function (*E*_*det,K*_ (*m*) = (*mG*^*′*^ + *rK*)), where *G*^*′*^ is a base point of the curve, possibly different from *G*. In our distributed scenario, it is possible to split the value *r* and the base point *G*^*′*^ among the computing nodes (analogously to the way the collective public key is split), and run an interactive protocol that converts a probabilistic encryption of a message into a deterministic one. Further details about this protocol can be found in (*33*).

It must be noted that, due to its equality-preserving property, deterministic encryption is susceptible to frequency attacks. That is, any attacker that knows the frequency distribution of the clear-text terms can try to map them to the frequency distribution of the corresponding encryptions, hence braking it. In this work, we rely mainly on probabilistic homomorphic encryption, and whenever we need deterministic encryption, we apply additional countermeasures (dummy generation, see Section 2.4) to avoid this kind of attacks.

#### 1.2 Integrity, Accountability and Access Control

There are two main approaches to protect data integrity and enforce access control in an accountable way: centralized and distributed. We briefly introduce the advantages and disadvantages of each of them, justifying our choice of a distributed approach for our system.

##### 1.2.1 Centralized Approach

Enforcing access control has been traditionally the job of centralized (cloud) services (*34*). These implement the logic that stores and interprets the rules applied to grant or deny access to the sensitive data, and also log the access requests and their outcomes. These services are typically manually configured and vulnerable to compromises. In personalized medicine, however, due to the sensitivity of the managed data and the inherently distributed nature of the data sources, relying on the existence and good will of such a provider or centralized authority poses significant risks that the multiple data holders are rarely willing to accept, especially when considering networks of clinical institutions or large sets of individuals. The single point of failure represented by the centralized authority becomes an obvious target for attackers, and represents a threat for the data in the whole network if compromised. Furthermore, the recent growth of the personal genomics industry has exacerbated existing privacy concerns due to widely publicized security breaches, nontransparent data sharing practices and government access to genomic data without consent.

##### 1.2.2 Distributed Approach

In a distributed environment, a centralized authority is avoided, and accountability usually relies on trusting each and every of the stakeholders involved in the collaborative processes (federated network). Our approach prevents this weakness by relying on a distributed ledger technologies (DLTs, a.k.a. blockchains) (*35–37*). DLTs are considered a core building block for many next-generation technologies relevant in multiple sectors of our future society, such as finance (*38*), healthcare (*39*) and e-democracy (*40*). The idea around distributed ledger is to move from a trust-and-hope model, where a centralized authority provides service to clients, to a trust-but-verify setting where the abstraction of this centralized authority is implemented by a (sub)set of the participants, who replicate the actions of the authority and collectively agree (by majority) on the ground truth. This split of trust makes the subversion of the system a challenging task for an adversary, as he would need to stealthily compromise the majority of participants, instead of a single central authority, in order to change the *ground truth*.

##### 1.2.3 Permissioned vs unpermissioned blockchains

There are two types of blockchains that can be deployed in a distributed scenario: unpermissioned and permissioned. The former are typically related to cryptocurrencies (e.g., Bitcoin), where there is no restriction to the entities that can join the management of the chain and modify its contents; the rules for updating the chain are normally based on *Proof of Work*, requiring that the participating entities (*miners*) perform some computationally costly operations before they are allowed to modify the chain by adding a new block. Besides not being environmentally friendly, these chains are susceptible to the so-called *51% attack*, in which a (minority) group of miners temporarily accumulate enough computational power to be able to drift the evolution of the chain at their will.

Contrarily, permissioned blockchains restrict the set of entities that have write-access to the chain, therefore effectively limiting the blockchain maintainers to a set of authorized parties. The evolution of the chain is not based on *proof-of-work*, but on strict consensus rules that define the majority ratio needed to add new blocks to the chain. Therefore, permissioned blockchains are a perfect fit for distributing the trust among a set of authorized servers, out of which the users only need to trust a majority, hence effectively avoiding single points of failure.

#### 1.3 Secure Data Release

Once data has been processed and is ready to be publicly released or shared with authorized entities, it has to be protected to avoid inference attacks that can single out individuals whose data has been used to produce the released results. Traditionally, statistical approaches are used to protect released data against these attacks, by means of modifications to the data that reduce its accuracy but limit the leakage that could lead to successful inferences. These approaches comprise de-identification and anonymization techniques for released individual data records, and differential privacy for released aggregated data.

##### 1.3.1 Data De-Identification

A data record is de-identified when direct identifiers are removed (*41*). Following the Privacy Rule of the Health Insurance Portability and Accountability Act (HIPAA), de-identification process can be applied following the Safe Harbor rule (removal of 18 personal identifiers), or by expert determination. In either of both cases, after the removal of direct identifiers it may still be possible to re-identify the individual in the presence of unique combinations of indirect identifiers, such as genomic data; therefore, de-identified records cannot be considered anonymized.

##### 1.3.2 Data anonymization

Data anonymization (*11*) is the result of processing personal data with the aim of irreversibly preventing re-identification of the data subject. Both direct and indirect identifiers are subjected to aggregation, generalization, suppression and/or randomization to mitigate re-identification risks. There are several metrics to determine the achieved degree of anonymization; one of the most popular is k-anonymity (*11*): A dataset fulfills *k*-anonymity whenever it contains at least *k* individuals with any chosen combination of quasi-identifiers. Therefore, each individual in the dataset is “hidden” within an anonymity set of at least *k* people.

##### 1.3.3 Differential Privacy

A function applied over a dataset is called differentially private (*12*) whenever the (distribution of the) result of such function does not substantially change by the presence or absence of one individual in the dataset. Therefore, the release of a differentially-private result reveals negligible information about each of the individuals in the dataset. Differential privacy is usually achieved through randomization techniques; i.e., adding a controlled amount of random noise to the output of the to-be-computed deterministic “exact” function *f* (·), such that the variance of the noise exceeds the maximum deviation of the function’s output produced by the contribution of any individual as part of the input dataset. There are several notions of differential privacy; the one we use in this work is (*ϵ, δ*)-differential privacy (*12*). Formally, this notion is defined in the following way: Given two databases *X* and *X* ^*′*^ that differ in one record, a randomized mechanism *κ* is said to be (*ϵ, δ*)-differentially private if for all possible outputs *y* (query responses) of *K*, it holds that *Pr*[*κ*(*X*) = *y*] *≤ e* ^*ϵ*^ *· Pr*[*κ*(*X* ^*′*^) = *y*] + *δ*, where *Pf* [·] represents the probability of an event. The informal interpretation of the previous expression is that the fact that two databases differ in one record does not significantly change the probability distribution of the output of the query. In practical terms, *κ*(.) is usually defined as the deterministic (exact) output of the desired query *f* (.) plus noise with a bounded power, usually drawn from a Laplacian or Gaussian distribution.

##### 1.3.4 Protecting Genomic Data Release

While anonymization and de-identification techniques can, to some extent, mitigate the reidentification risk of clinical data, genomic databases have been proven vulnerable against several types of inference attacks. Membership inference has been popularized by Homer et al. back in 2008 (*10*). It consists in inferring whether some target individuals contributed their genomic data to a cohort while knowing only part of the target’s data and summary statistics about the cohort. Another inference attack, variant inference (proposed by Humbert et al. (*42*)), can be carried out by relying on the intra-genome (between variants) and inter-genome (between relatives) correlations. Gymrek et al. have further shown that the identity of individuals could be inferred by leveraging genetic genealogy databases (containing surnames) with short tandem repeats on the Y chromosome (*3*). Shringarpure and Bustamante have also shown that it is possible to infer whether an individual is part of a group of interest by only asking binary questions about the presence or absence of alleles at different positions in the genome (*43*). It is worth noting that membership inference attacks have also been recently performed with relatively high success against genomic databases (e.g., Backes et al. (*44*)).

Consequently, genomic data cannot be effectively de-identified or anonymized, and it requires more strict protection mechanisms, such as encryption, to avoid re-identification risks.

#### 1.4 Related Work

Multiple solutions have been proposed in order to enable queries on a database while protecting the privacy and/or the confidentiality of the data provider. These solutions also aim at protecting the privacy of each individual storing its data in these shared databases. This can be achieved by relying on secure multi-party computation (*45*) or on differential privacy (*46*). Settings with a central database comprise the system by Hamlin et al. (*47*), who propose a solution based on multi-key searchable encryption that enables keyword-search on a database of encrypted terms, but it imposes a considerable storage overhead on the server by replicating each file every time it is shared. Popa et al. (*48*) propose an encrypted database enabling SQL queries over encrypted data, by relying on different forms of encryption depending on the allowed SQL function; nevertheless, it has been shown susceptible to a series of attacks, including frequency attacks on the deterministically encrypted terms in the database; i.e., an attacker knowing the distribution of the clear-text terms can revert the encryption. Finally, Meng et al. (*49*) propose a system that securely processes top-k ranking queries over outsourced encrypted databases. All these solutions are based on centralized databases with centralized processing, and hence they present a single point of failure that poses a threat for the stored data if either an attacker breaches the server or if it breaches the client storage to get access to the key.

Commercial systems targeting the protection of database contents have been produced by Microsoft (Always Encrypted database engine), which relies on deterministic encryption on a cloud-hosted database and off-cloud key-stores used by the client to build the queries. Unfortunately, it can be susceptible to frequency attacks if the distribution of the clear-text query-terms is known, and the cloud database is again a single point of failure. Distributed commercial systems comprise Cybernetica’s ShareMind (*50*) and Inpher’s XOR (*51*), both targeted at the execution of machine learning functions on data split among a set of computing servers, relying on secret-sharing and secure multiparty computation techniques.

In order to compare genomic data among different patients in an honest-but-curious model, Jagadeesh et al. (*5*) propose secure two-party computation method such that patients encode their genomic variants and the system ensures that a query outputs only some statistical (max, setdiff,…) information about the queried variants and nothing else about the patients. On the other hand, a system (*33*) combining homomorphic encryption and differential privacy in a decentralized way (UnLynx), was proposed to enable statistical queries on distributed databases while avoiding single points of failure for achieving the sought security and privacy guarantees; this system was later on instantiated in a medical scenario in MedCo (*52*), enabling privacy-preserving queries to distributed databases of clinical and genomic data.

It is worth noting that none of these systems features the citizen-centric approach that we propose, as they do not address access control, consent management, or accountability requirements sought by individuals. Moreover, they do not consider individual citizens as data providers but they mostly address hospitals and research institutions’ privacy and security concerns.

### 2 Materials and Methods

#### 2.1 System and Threat Models

As opposed to prior work focusing on institutional-based data provision, the system we propose enables for the first time individual citizens to directly share their clinical and genomic data with researchers without compromising their privacy and by keeping control on who can access and use their data. We model such system with the following four parties (see Figure S1):

- **Data Providers (DP)** are individual citizens who use the platform to securely share their clinical and genetic data for population genomic studies.
- **Data Queriers (DQ)** are researchers who use the platform to find individuals with interesting clinical and genetic characteristics in order to recruit them in clinical research studies or pharmaceutical trials.
- **Storage Units (SN)** can host of one or multiple servers and they are responsible for securely storing the clinical and genomic data of data providers. Data providers can choose any of the available storage units for storing their data. Such units can be represented by any governmental, academic or commercial institution providing an IT infrastructure capable of storing large volumes of data, e.g., a public cloud provider such as Amazon Web Services or Google Cloud.
- **Computing Nodes (CN)** are a set of independent governmental, academic or commercial institutions hosting one or several servers that are responsible for jointly and securely processing data discovery requests from data queriers.

According to the definitions of possible adversarial behaviors reported in Box 1, we assume that *storage units* and *computing nodes* are honest-but-curious (HBC), non-colluding parties.

**Figure S1:**
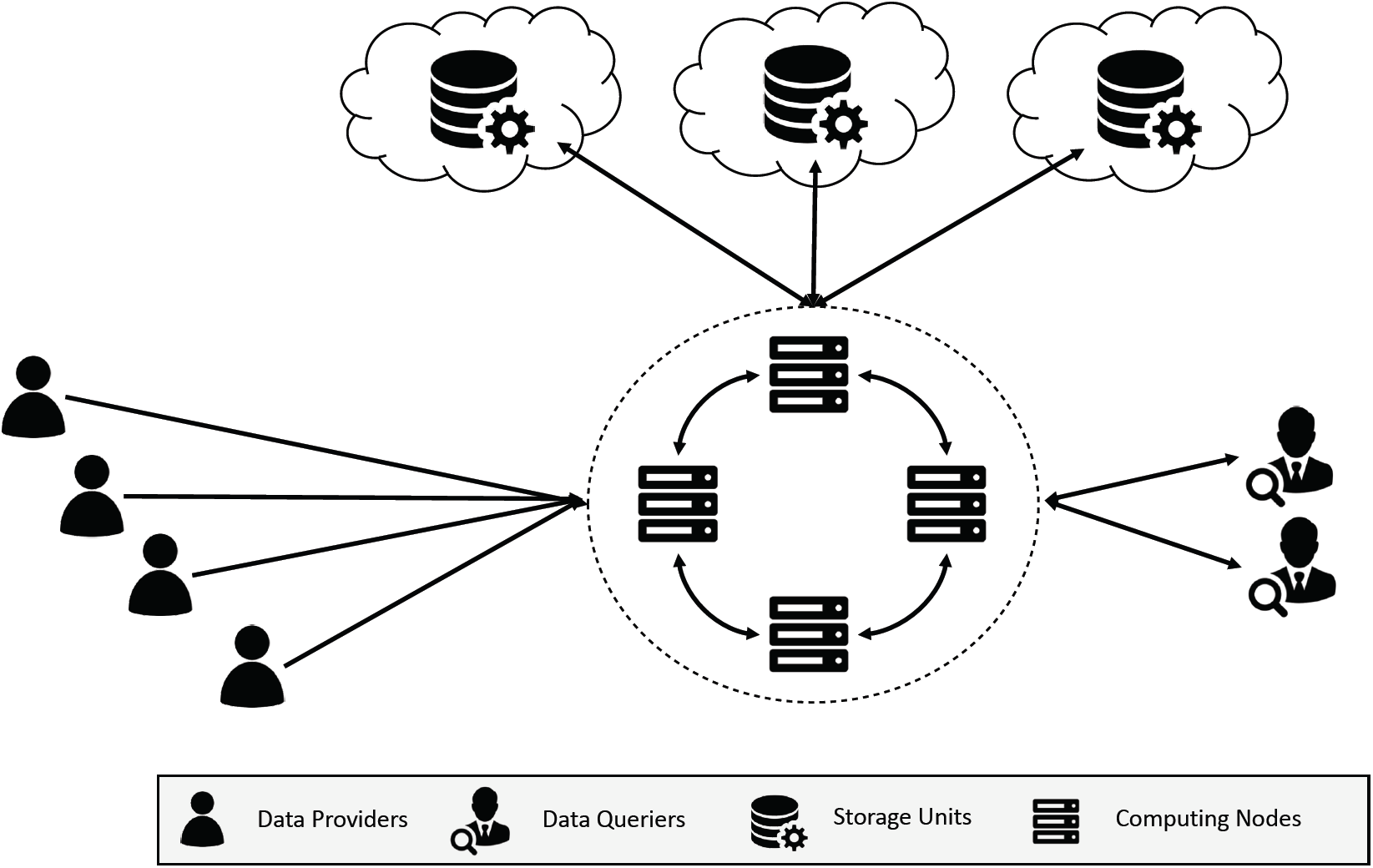
High-level representation of the system model.

This means that they honestly follow the protocol but might try to infer sensitive information about data providers and data queriers from information processed or stored at their premises during the protocol. The HBC adversarial model is typically used in information security to model the behavior of storage and computing IT service providers such as public clouds. Cloud providers have business incentives for not colluding with competitors and for not tampering with their customers’ data and operations. Yet, they can still be “curious” and (if compromised by external (hackers) or internal (insiders) attackers) gather or infer sensitive and private information about their customers stored or processed at their premises.

We assume that data providers are honest (H), i.e, they only expose their real and accurate clinical and genomic data without trying to fool the system with fake information. We acknowledge that the proposed platform could potentially be abused by dishonest individuals who would lie to sound “more interesting” to pharmaceutical companies and for being recruited in clinical trials.

However, we note that it is very unlikely that this misbehavior occurs on a large scale. First, many researchers are also interested in recruiting a lot of control samples (not only case samples); this means that there is no obvious way to make a clinical and genetic profile more “appealing” for clinical trials. Second, once the data queriers get access to the raw data, they will be able to request the data providers other samples for further genetic analyses or to get access to their electronic medical record, thus unmasking potential liars.

Finally, we consider data queriers as potentially misbehaving parties, as their credentials can be stolen by malicious attackers who can use the system in an unethical way in order to obtain sensitive information about data providers. In other words, an attacker impersonating a legitimate user of the system can potentially perform specific queries to try to infer undisclosed sensitive attributes about a targeted individual, e.g., membership to a sensitive cohort (*10, 43*), or reconstructing the individual profile of as many data providers as possible by performing an unlimited number of requests.

##### BOX 1

###### Definition of adversarial behaviors

- Honest (H): a honest party honestly follows the protocol without trying to obtain sensitive private information about other parties
- Honest-but-curious (HBC): a HBC party honestly follows the protocol but might try to obtain sensitive private information from other parties’ data stored or processed at its premises
- Malicious (M): a malicious party might tamper with the protocol/data in order to obtain sensitive private information about other parties or/and influence the output of the protocol for its personal advantage or profit

#### 2.2 Data Encoding

The proposed system enables data providers to securely both their clinical and genomic data. In the following, we describe how these different data types are encoded and stored within our platform.

##### 2.2.1 Genomic Data Encoding

We consider genomic variants regardless of the format in which they are initially called after sequencing. A variant is uniquely identified by its chromosomal position (CHR, POS), the reference allele (REF) and the alternate allele (ALT). In general, this information is public and considered to be non-sensitive meta-data. What is sensitive and must be protected is the association of variant meta-data an individual genotype, i.e., the value that the variant takes for a given individual. In particular, we consider that an individual variant genotype is sensitive when it is “mutated”, i.e., if there is at least one of the two alleles composing the genotype that has changed from the reference to the alternate allele. Therefore, individual variants genotype can be encoded as strings of the form “CHR:POS:REF>ALT” which means that, at the given chromosomal position (CHR:POS), the individual carries at least one allele that has mutated from the reference to the alternate (REF>ALT). In order to encrypt such string, we transform it into a 64-bit integer with the following convention: 1 bit indicating that the code encodes a genetic variant, 5 bits for the chromosome identifier, 28 bits for the position, 15 bits for the reference allele and 15 bits for the alternate allele.

##### 2.2.2 Clinical Data Encoding

In this work, we consider self-reported clinical data. This can include diagnoses, medications, procedures, simple findings and demographics information. We refer to each clinical data point as “clinical attribute”. In general, clinical attributes can be encoded by using standard medical ontologies and terminologies such as SNOMED-CT (*53*), ICD10 (*54*), or RxNorm (*55*). In such ontologies, clinical attributes are represented by “concepts” that are uniquely identified by an alphanumeric code. For example, “lung cancer” is encoded by the code “C34” in the ICD10 terminology. As for the genomic encoding, we convert alphanumeric concept codes into 64-bit integers.

##### 2.2.3 Data Model

In order to effectively store and perform data discovery queries based on Boolean combination of genomic variants and clinical attributes on data of potentially millions of individuals, we rely on the well-known “star-schema” relational data model typical of clinical data ware-houses. The star-schema data model is based on the Entity-Attribute-Value (EAV) concept and it is adopted by several state-of-the-art clinical data discovery systems such as i2b2 (*56*) and tranSMART (*57*).

Figure S2 shows a toy example for two data providers and illustrates the main principles behind this data model. Clinical attributes and genetic variants represent “observations” about individuals and are stored in a narrow central table called “observation fact”. In this table, each row stores the code of one observation uploaded at a given timestamp by a given data provider. The other tables called “dimensions” store the meta-data about data providers and the clinical attributes and genetic variants uploaded to the system. For example, the “data providers” dimension stores meta-data about data providers such as their pseudonymized demographic information (pseudoidentifier, ethnicity, gender, age, etc.), whereas the “concept” dimension stores meta-information about the uploaded observations, i.e., the ontology codes used to encode self-reported clinical variables and the genetic codes used to encode genetic variants.

From a data protection perspective, the information in the model to be protected from potential adversaries is represented by the link between the data providers and their clinical attributes and genomic variants stored in the “observation fact” table. In the following, we demonstrate how we can protect such link by encrypting concept codes and making them indistinguishable all the while enabling efficient data discovery queries.

**Figure S2:**
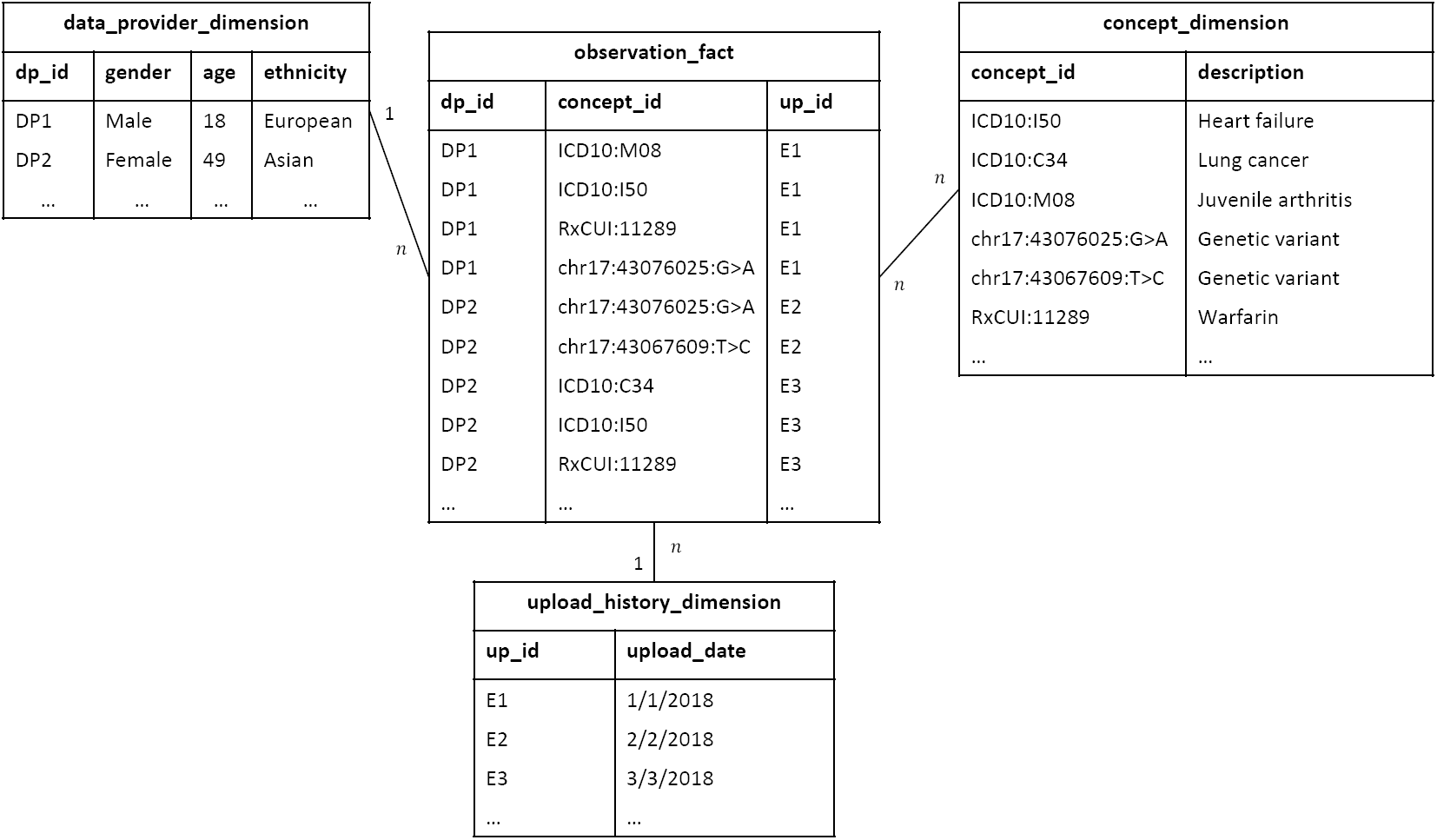
Toy example of the “star-schema” data model storing data uploaded by two data providers DP1 and DP2. The *observation fact* contains clinical and genetic observations for the two data providers (one observation per row). The meta-data about data providers is stored in the *data provider dimension* table. The *concept dimension* table stores information about clinical and genetic variables (or concepts). The *upload history dimension* table stores meta-information about the data upload, e.g., the data upload date.

#### 2.3 Privacy-Preserving and Secure Data Sharing Protocol

In this section, we provide a detailed description of the privacy-preserving and secure protocols that operate our platform and enable researchers to (i) efficiently run data discovery queries on encrypted data and (ii) securely access consented individual-level records, in case more in-depth analyses are needed.

**Figure S3:**
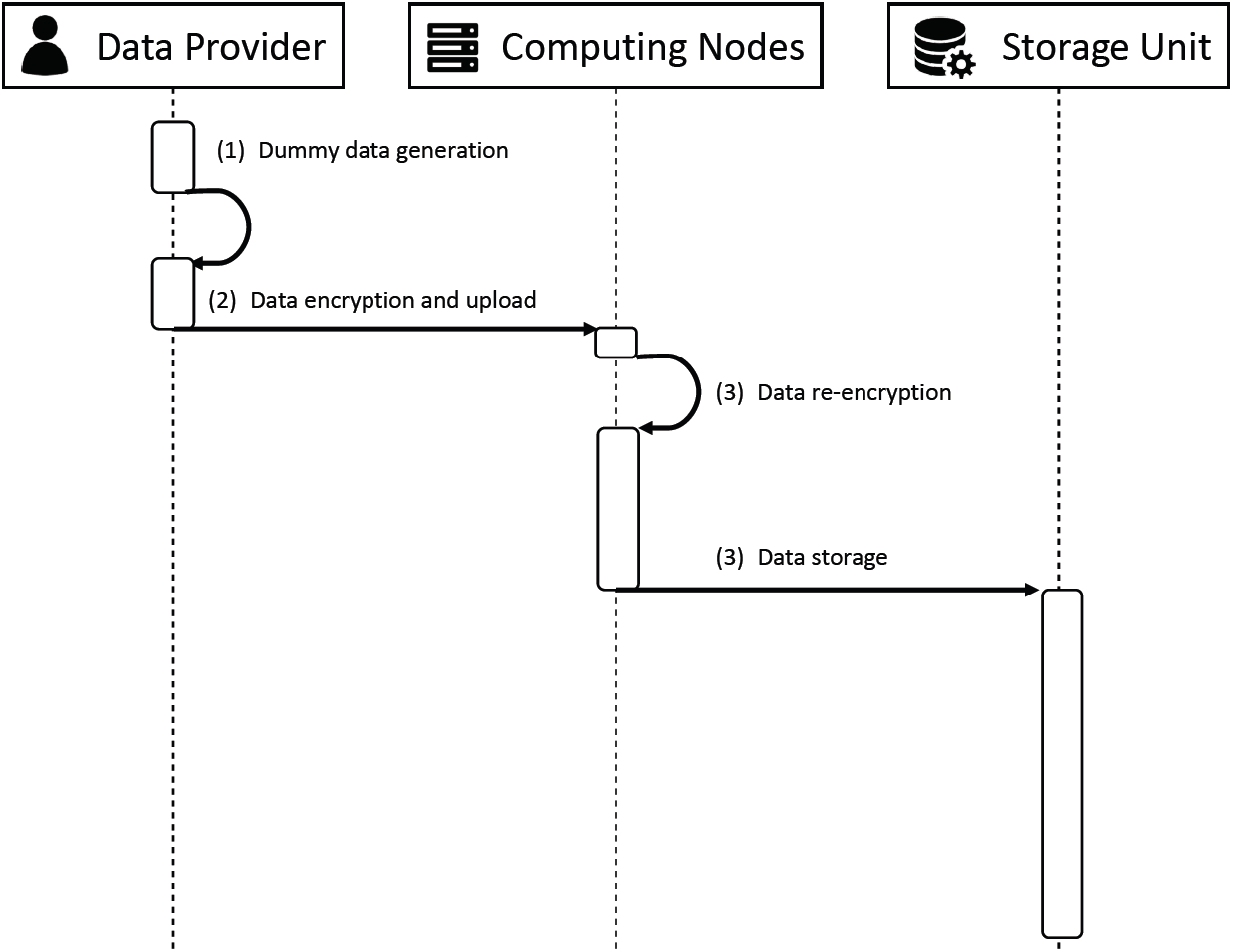
Data preparation phase sequence diagram.

After the platform initialization (performed only once), we distinguish three sequential phases: a data preparation phase that is performed by each data provider each time clinical and genetic variables are uploaded to the platform, a data discovery phase, and a data access phase that are performed each time a researcher wants to run data discovery queries or request access to individual-level records.

##### 2.3.1 Platform Initialization

Let **I** designate the set of independent but collaborative computing nodes that operate the platform. During the initialization phase, each computing node (*i* ∈ **I**) generates a pair of public/private keys (*k*_*i*_,*K*_*i*_) for the EC-ElGamal additively-homomorphic encryption scheme along with a secret key *s*_*i*_. Then, all computing nodes combine their public keys in order to generate a single public key *K* = ∑_*i*_ *K*_*i*_ that is used by data providers to encrypt their data before uploading them to the platform. We denote this key as the *collective encryption key*.

This “joint” key generation technique ensures that data encrypted under the collective key is protected unless all computing nodes are compromised and their individual private keys stolen. The more computing nodes participate in the generation of the collective key, the higher is the overall security of the system. Finally, along with the key generation, the computing nodes set up a private permissioned blockchain to be used as auditable and immutable log for the operations executed on the platform by the different parties and for storing topology information about storage units.

##### 2.3.2 Data Preparation Phase

The data preparation phase consists of four steps, also represented by the sequence diagram in Figure S3. For the sake of the presentation and without loss of generality, we describe the different steps of this phase by considering only a single data provider and a single storage unit.

###### (1) Dummy data generation

The data preparation phase starts with a dummy data generation algorithm. This algorithm, explained in detail in Section 2.4, takes as input the set of clinical attributes and genetic variants, also referred to as “observations”, of the *j*-th data provider (**V**_**j**_) along with the corresponding population statistics (i.e., the frequency distribution of genetic variants and prevalence of clinical variables), and outputs a set of dummy individuals with a plausible set of observations specifically selected to flatten the global joint distribution of observations. We denote as 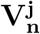 the set of observations for the *n*-th dummy individual in the set **N**^**j**^ generated from the *j*-th real individual. To prevent that the fake observations affect the data discovery process, a binary flag is assigned to each individual and appended to the corresponding set of observations. Such flag is set to 1 for real individuals and to 0 for dummy individuals.

###### (2) Data encryption and upload

After the dummy data generation, the data provider generates a symmetric encryption key (*S*_*j*_) and uses it to encrypt the original files containing her genetic variants and self-reported clinical attributes (e.g., VCF files and PDF clinical surveys/reports). Then, the data provider uses the collective encryption key *K* to encrypt:

- the symmetric encryption key as *E*_*K*_(*S*_*j*_),
- the codes of her observations and those from her dummies’ 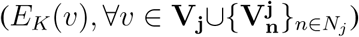,
- the set of binary flags (*E*_*K*_(*f*_*j*_) and *{E*_*K*_(*f*_*n*_)*}*_*n*∈*N*_*j*).

The data upload generates a blockchain transaction which is verified by all computing nodes and immutably stored on the blockchain. The transaction contains the data access policy and the mapping between the data provider’s pseudo identity and the set of pointers to the uploaded encrypted files storing the genetic and clinical variables.

###### (3) Data re-encryption

The computing node receiving the data initiates a distributed re-encryption protocol on the encrypted observation codes. This protocol switches the encryption of the codes from homomorphic to equality-preserving, thus enabling the execution of equality-matching queries on the re-encrypted codes. Homomorphically encrypted flags and symmetric encrypted files are not involved in this protocol, and are sent for storage to the storage unit. As explained in Section 1.1.3, the use of equality-preserving encryption reveals the distribution of the encrypted observation codes. However, thanks to the presence of the previously generated dummy individuals and their encrypted observation codes, it is impossible for the adversary to perform frequency attacks, as the overall distribution of encrypted codes is uniform, and codes are indistinguishable from each other. This protection is guaranteed as long as the binary flags remain homomorphically encrypted and dummy individuals cannot be told apart from real individuals. Let *E*_*K*_(*υ*) = (*C*_1_, *C*_2_) = (*rG, υ* + *rK*) be the expanded representation of an EC-ElGamal encrypted observation code *υ* under the collective public key *K*, where *r* is a random nonce and *G* is the base point of the elliptic curve. The distributed re-encryption protocol consists of two rounds that involve all computing nodes operating the system. In the first round, each computing node *i* sequentially uses its secret *s*_*i*_ and adds *s*_*i*_*G* to *C*_2_. After this first round, the resulting ciphertext is 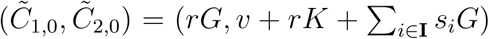. In the second round, each computing node partially and sequentially modifies this ciphertext. More specifically, when the *i*-th computing node receives the modified ciphertext 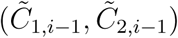 from (*i* − 1)-th computing node, it computes 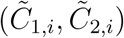, where 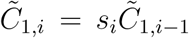 and 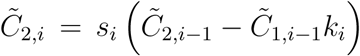. At the end of the second round, the equality-preserving version of the encrypted variable code is obtained by keeping only the second component of the resulting ciphertext 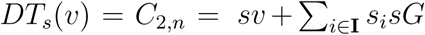, where *s* =П_*i*_*∈***I** *s*_*i*_ is the collective secret corresponding to the product of each computing node’s secret that was generated during the initialization phase.

###### (4) Data storage

Once the re-encryption of clinical and genetic variables is terminated, the computing node who initiated the protocol sends them to the storage unit for storage under the star-schema data model. Figure S4 shows the same toy example of Figure S2 with the encrypted data an the addition of a dummy individual necessary to flatten the distribution of encrypted codes.

##### Data Discovery Phase

The data discovery phase consists of six steps also represented by the sequence diagram in Figure S5. Similarly to the previous phase, for the sake of the presentation and without loss of generality, we describe the different steps of this phase by considering only a single data querier and a single storage unit.

###### (1) Query Generation

Upon registration to the platform, a data querier *q* is assigned a query budget *∈*_*q*_ that is stored on the blockchain and consumed each time he sends a new queryn to the system. The goal of this query budget is to limit the total number of queries that can be sent by the same data querier and, as a consequence, the amount of undisclosed sensitive information about data providers that could potentially be inferred from query results. After proper authentication, an authorized data querier who has enough query budget can run a secure data discovery query and, depending on his authorization level, obtain either the total number of individuals on the platform whose data matches the set of inclusion/exclusion clinical and genetic criteria specified in the query (also referred to as query items) or the identifiers of these individuals. As such, the data querier generates his own pair of EC-ElGamal public/private keys (*U*_*q*_,*u*_*q*_) and builds a query by logically combining (i.e., through Boolean AND and OR operators) clinical and genomic codes from the list of all possible codes present on the system. Then, the selected codes are encrypted with the collective encryption key *K* and sent along with the data querier’s public key *U* to one of the computing nodes. As for the data upload, also the query upload generates a blockchain transaction that gets verified by all computing nodes of the platform and immutably stored on the blockchain. The query transaction contains the query definition and the pseudo identity of data querier.

**Figure S4:**
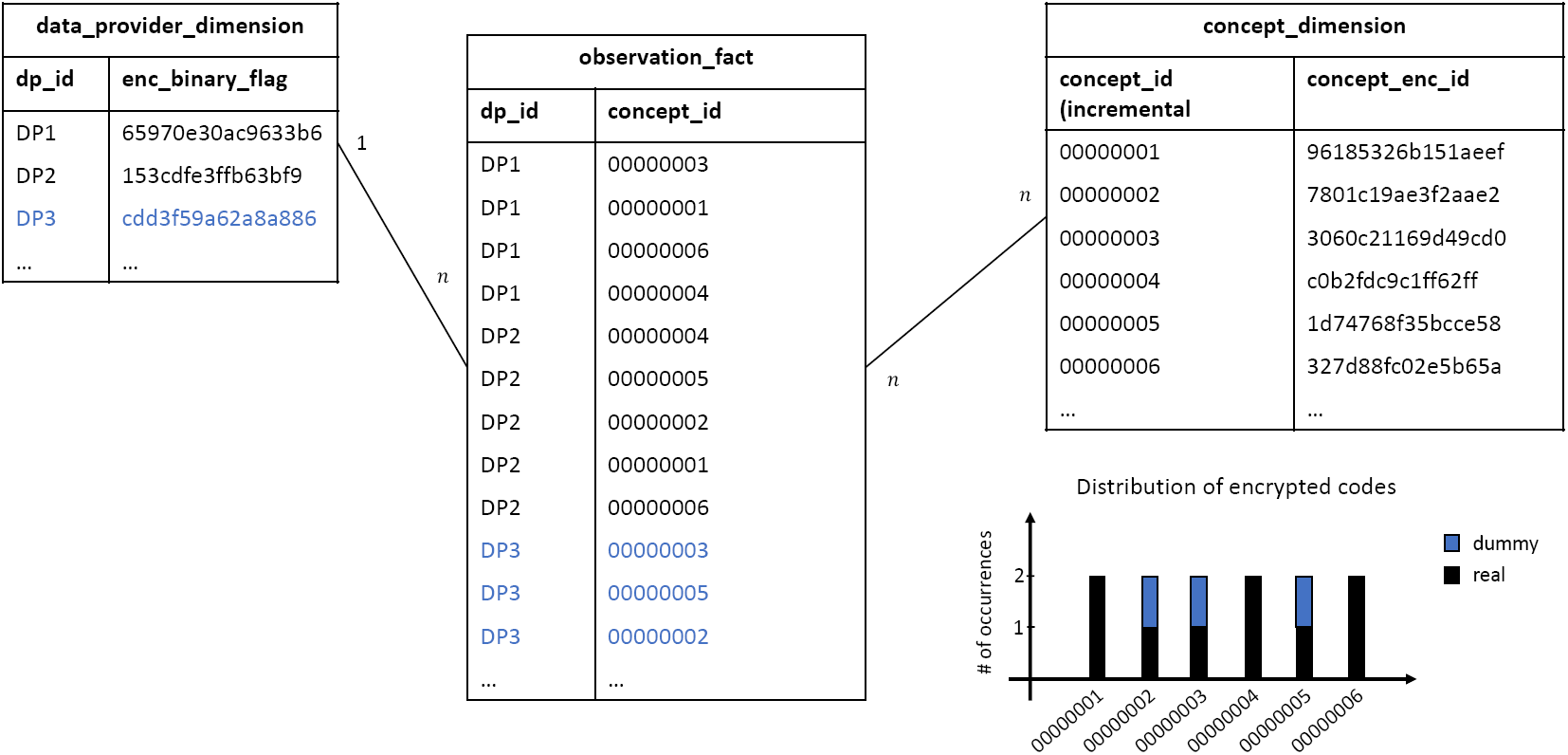
Toy example of Figure S2 with only the tables of the “star-schema” data model affected by data encryption. A dummy individual (DP3) with three observations has been added in order to flatten the distribution of encrypted observation codes.

**Figure S5:**
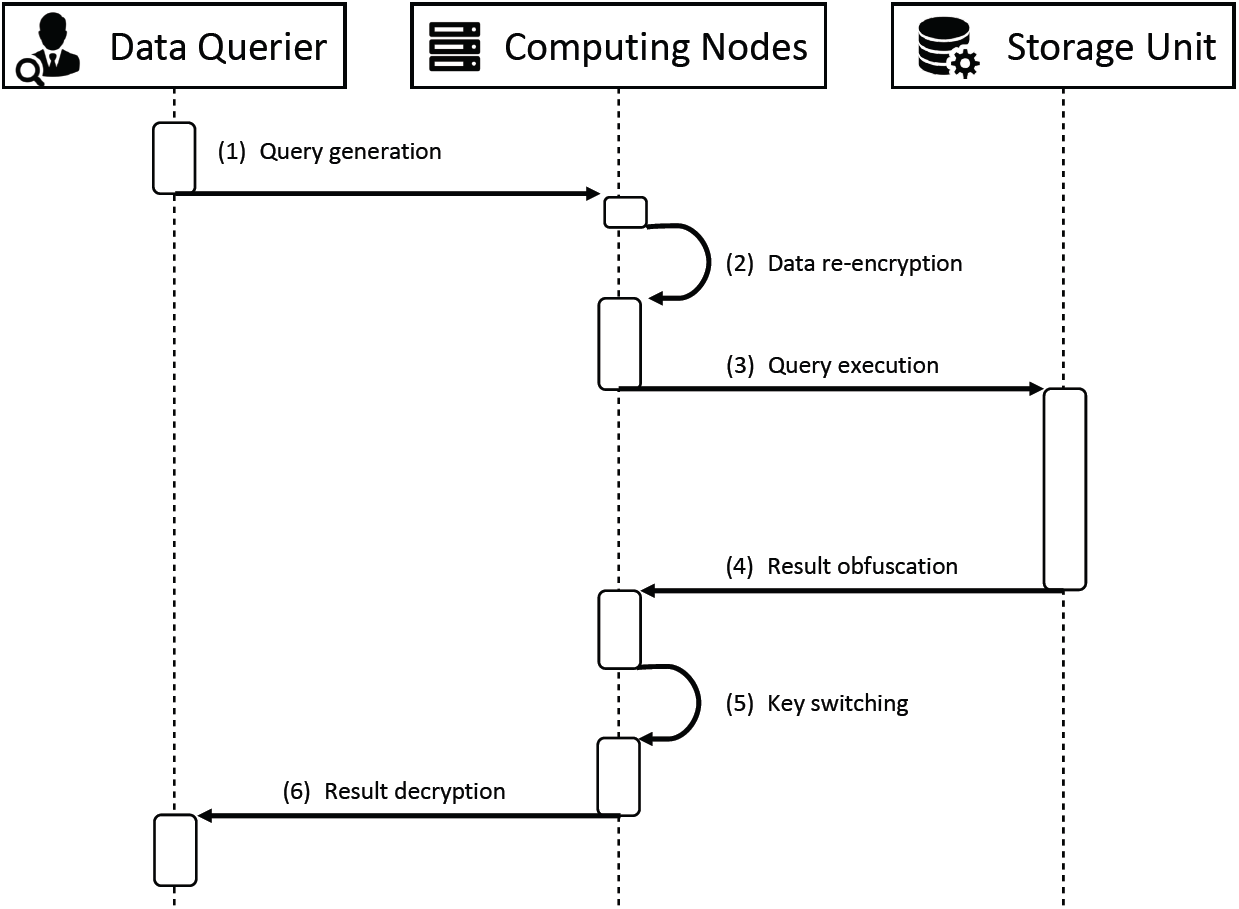
Data discovery phase sequence diagram.

###### (2) Query re-encryption

In order to match the codes of the observations uploaded by the data providers, the codes used in the query are also re-encrypted from the initial homomorphic form to the equality-preserving form. To this purpose, the computing node receiving the query initiate the same distributed re-encryption algorithm described in step (3) of the data preparation phase and, once the protocol is over, it forwards the query with the re-encrypted codes to the storage unit.

###### (3) Query execution

The storage unit executes the query and retrieves the encrypted binary flags for the individual who have encrypted observations codes that match the ones in the query. Let **Φ** be the set of identifiers of individuals (both real and dummy) selected by the query, if the query requests the size of this set, the flags are homomorphically summed as *E*_*K*_(*R*) = *E*_*K*_(∑_*j*_,*n∈* **Φ** *f*_*j,n*_) = *j,n∈* **Φ** *E*_*K*_(*f*_*j,n*_) and the encrypted result is returned to the computing node who sent the query. Note that in this case, the dummy individuals fetched by the query are automatically filtered out from the result as their encrypted binary flags sum to the encryption of zero *E*_*K*_(∑ _*n∈***Φ**_ *f*_*n*_) = ∑ _*n∈***Φ**_ *E*_*K*_(*f*_*n*_) = *E*_*K*_(0). If the query requests the identifiers of the individuals selected by the query, the latter are returned after having been masked by a homomorphic scalar multiplication with the corresponding encrypted flag, *E*_*K*_(*f*_*j,n*_) *× id*_*j,n*_*∀j, n ∈* **Φ**. The masking is necessary for concealing the identifiers of the dummy individuals to the data querier as each of them will decrypt to zero. In both cases, the query execution generates a blockchain transaction that contains the query definition and the identifiers of the selected individuals and is immutably stored on the blockchain after verification.

###### (4) Result obfuscation

This steps only takes place if the query requests the size of the set of individuals matching the query. Depending on the authorization level of the data querier, computing nodes can run a distributed obfuscation protocol so that the released result cannot be used to infer further sensitive information about data providers. This protocol, firstly introduced in a previous work (*33*) in which some of the co-authors of this paper were involved, enables a set of computing nodes to collectively and homomorphically add to the query result a noise value, sampled from a probability distribution that satisfies the differential privacy requirements. In this way, (*ϵ, δ*)-differential privacy is ensured for each data provider without revealing to any entity the amount of noise added. The protocol operates in two phases: an initialization phase, executed during the platform setup, and a runtime phase executed for each new query. In the initialization phase, a randomly selected computing node generates a probability distribution curve (e.g., Laplacian distribution or Gaussian distribution) based on the publicly defined differential privacy parameter (*ϵ*). The same computing node samples a list of integer noise values that approximate such distribution and stores the list along with the distribution parameters on the system blockchain. In this way, every party involved in the protocol can verify the correctness of the sampling procedure. In the runtime phase, the computing node that receives the encrypted query results from the storage unit fetches the list of sampled noise values from the blockchain, encrypts them with the collective public key *K* and starts a distributed secure shuffling sub-protocol on this list. This sub-protocol is based on the original idea proposed by the Neff shuffle algorithm (*58*) and enables computing nodes to sequentially shuffle a list of encrypted values so that the final list cannot be linked to the original one by any node. More specifically, the protocol takes as input a list of EC-ElGamal encrypted values (*C*_1,*i,j*_, *C*_2,*i,j*_) forming a *a × b* matrix, and outputs a shuffled matrix of 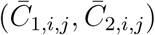 pairs such that for all 1 ≤ *i* ≤ *a* and 1 ≤ *j* ≤ *b*, 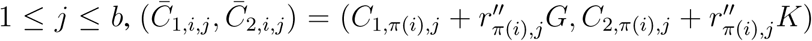, where 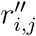 is a re-randomization factor, *π* is a permutation, and *G* is the elliptic curve base point. Once the distributed secure shuffling is over, the same computing node who initiated the sub-protocol selects the first element in the list and homomorphically add it to the encrypted query result thus obtaining 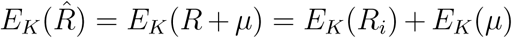, where *E*_*K*_(*µ*) is the encrypted noise value. Finally, the query budget of the data querier (*q*) is updated (*ϵ*_*q*_ −*ϵ*) and stored on the blockchain along with the used encrypted noise value. If the same query is asked multiple times by the same data querier (i.e., if the set of individuals’ identifiers resulting from two queries is the same), the querier’s budget is not reduced and the noise of the previous query is re-used to ensure that it cannot be averaged out in case of repeated equivalent queries.

###### (5) Key Switching

In order for data querier to decrypt the query result, the encryption the latter (total number of matching individuals or their encrypted flags) need to switched from the collective public key to the data querier’s public key. Computing nodes jointly run a distributed key switching protocol that enables them to perform this operation without ever decrypting the data. Let *E*_*K*_(*R*) = (*C*_1_, *C*_2_) = (*rG, R* + *rK*) be the EC-ElGamal encryption of the query result with the collective public key *K* and *U* be the data querier’s public key. The protocol starts with a modified ciphertext tuple 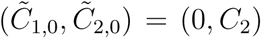. Then, each computing node partially and sequentially modifies this tuple by generating a fresh random nonce *v*_*i*_ and computing 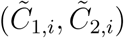 where 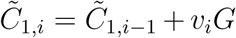 and 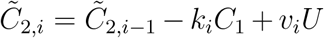. The resulting ciphertext corresponds to the query result *R* encrypted under the public key 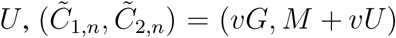 from the original ciphertext (*C*_1_, *C*_2_), where *υ* = *υ*_1_ + … + *υ*_*n*_. Finally, the key-switched encrypted result is sent back to the data querier.

###### (6) Result Decryption

The data querier uses his private key *u* to decrypt the query results. After decrypting the flags, the identifiers of the real individuals are separated from the ones of the dummy individuals.

##### 2.3.3 Data Access Phase

The data access phase enables the data querier to access the raw individual-level data necessary to perform analyses that go beyond the data discovery. This phase consists of five main steps also represented by the sequence diagram in Figure S6.

###### (1) Data Access Request

The data access phase starts with the data querier who requests access to a file containing raw clinical or genomic information (e.g., a VCF file). The data access request generates a transaction on the blockchain that contains the cryptographic hash-based signature of the file and information about the data querier such his name, affiliation and the description of the conducted study requiring access to individual-level data.

###### (2) Access Policy Verification

When data providers register on the system, they write an access policy that is stored on the blockchain. Two basic options are available: broad consent and dynamic consent. Raw files’ hash signatures and access policies are linked to a data provider identity (her public key) that is used to generate transactions for registration and data upload. Thus, by reading the blockchain, computation nodes can link data providers’ files to their correspondent access policies. If a broad consent is verified, the computing nodes proceed to make the data accessible. If a dynamic consent policy was chosen, the system front-end used by the data provider generates a notification when it synchronizes with the blockchain state. The data provider can review the information that is included in the data access request and then approve or reject the request by generating another transaction that is verified by the computing nodes.

**Figure S6:**
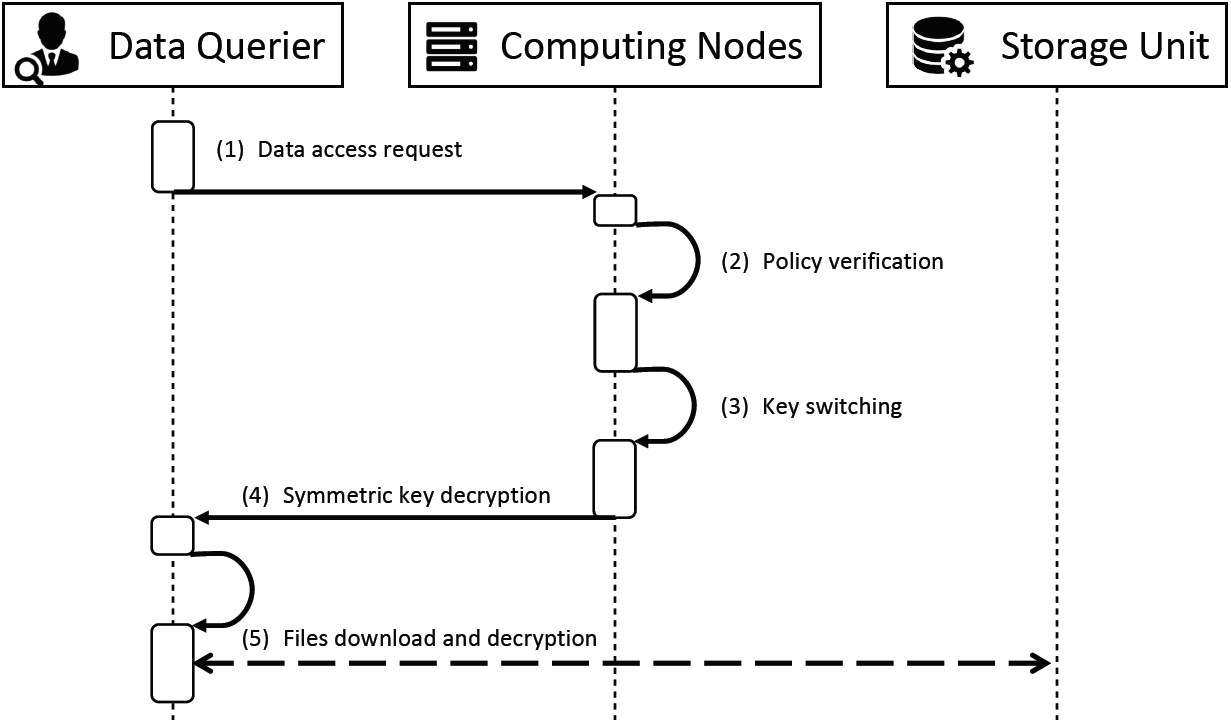
Data access phase sequence diagram.

###### (3) Key Switching

After access policy verification, the encryption of the symmetric key used to encrypt the raw files is switched from the collective public key to the querier’s public key. The process is the same as the key-switching at the end of the data discovery phase.

###### (4) Symmetric key decryption

The data querier decrypts the symmetric key with his private key

###### (5) Files download and decryption

Finally, the data querier uses the decrypted symmetric key to decrypt the raw files downloaded from the storage unit.

#### 2.4 Dummy Data Generation

In the proposed system, matching operations in the encrypted space between the codes of the uploaded observations and the codes in the query are possible thanks to the equality-preserving property of the encryption. During the re-encryption protocol described at step (3) of the data preparation phase, none of the computing nodes knows the secret used to re-encrypt the observation codes, and therefore, dictionary attacks are avoided as long as there is at least one honest node. Nevertheless, due to the characteristics of the equality-preserving encryption, frequency attacks are still possible if no further measure is taken. An attacker with knowledge of the population statistics would be able to map the histogram of the encrypted codes to the known frequencies of occurrence of each code, hence breaking the encryption. In order to avoid this type of attacks, we introduce an additional step during the data preparation phase for *generating dummy patients*. This step follows the same reasoning and analysis presented in our prior work (*52*), adapted to the patient-centric model proposed in this paper. We detail here the followed process and the main differences with respect to the process in (*52*).

The target of the dummy generation process is to maximize the confusion of an attacker with access to the encrypted real and dummy observations when trying to map encrypted observation codes with clear-text observation codes. As shown in (*52*), this can be achieved by flattening the empirical joint distribution of the observed codes, by producing dummies that “waterfill” their distribution. Unfortunately, this strategy produces a number of dummy individuals that grows exponentially with the number of observed codes, which is in general unfeasible. Several strategies can be employed to achieve a reasonable trade-off between privacy and number of dummy individuals. In particular, given *n* different codes observed in the database, (i) the codes can be clustered according to their source distribution (i.e., the frequency distribution of genetic variants and prevalence of clinical attributes), forming groups of similar-frequency codes with a minimum size of *n*^*′*^ *« n* codes per group, and (ii) the codes can be assumed to be independent across groups (i.e., the codes assigned to one block only influence other codes in the same group). In order to guarantee that the dummy individuals are indistinguishable from real ones, the former are generated with a set of codes that is a random permutation of the codes present in each real individual. This strategy minimizes the number of needed dummies while maximizing the anonymity set of codes, and therefore the attacker confusion.

When the whole database is known in advance by the data provider, it is possible to exactly adapt the empirical distribution to make it perfectly flat per cluster (as proposed in (*52*)). Contrarily, in our system model, each data provider is just one individual submitting one individual set of codes, without knowing the exact histogram of the other codes in the database. Hence, as the exact empirical distribution is not available, the system uses instead the publicly available frequency/prevalence for each of the available codes and applies the clustering based on this “theoretical” distribution. The cluster associations for each observation code and the target cluster-flat distribution are published and communicated to each data provider; the computation of these clusters and of the target distribution can also be published in the blockchain for auditability. When a data provider wants to submit an observation a set of codes to the system, it produces a fixed number of dummies as random permutations of the codes; these permuted records approximately follow the “reversed” population distribution (published by the system and auditable through the blockchain) that will guarantee that the union of dummy and real individuals will converge to a flat distribution. The reversed distribution can be obtained as the normalized distribution that, added to the original population probability density, will waterfill it to make it flat; the dummy ratio (number of dummies per real patient) can be adjusted as the weighting term between the real and the dummy distribution to make them sum up to a completely flat uniform distribution.

#### 2.5 System Implementation

We implemented the proposed system by combining different components of several open-source and widespread software platforms.

For the core back-end components (database and users management) we used the state-of-the-art open-source tool for clinical data discovery, Informatics for Integrating Biology and the Bedside (i2b2) (*56*). i2b2 is one of the most widespread tools to enable secondary use of electronic health records. It is currently used by more than 300 medical institutions world-wide to store patients’ clinical and genomic observations. i2b2 is implemented in Java and it is optimized for running data discovery queries on the star-schema data model described in Section 2.2.3.

For the secure privacy-preserving component, we rely on the MedCo open-source framework (*52*) implemented in Go, which itself relies on the distributed cryptographic library UnLynx (*33*). We extended the MedCo secure protocols by implementing from scratch the new dummy data generation algorithm described in Section 2.4, the retrieval and masking of individuals identifiers matching the query described in Section 2.3, and the distributed obfuscation protocols also described in Section 2.3. We integrated all these updates directly into the MedCo codebase.

For the blockchain part of our system, we implemented our own permissioned blockchain by using the open-source Exonum framework (*59*). The Exonum framework enables the implementation of high-performance, permissioned blockchains that offer transparency and security that is comparable to the one of public (permissionless) blockchains. In contrast to other permissioned blockchains, Exonum uses a byzantine fault-tolerant (BFT) consensus algorithm that protects not only against breakdowns of the computing nodes but also against malicious behaviors. The BFT algorithm is computationally intensive, but because Exonum is written in Rust, one of the fastest programming languages, high transaction throughputs are possible (see Figure S10 for performance benchmark of our implementation). Additional security mechanisms include cryptographic commitments using Merkle trees that allow clients to verify responses that they receive from permissioned nodes and anchoring of transaction logs in the Bitcoin blockchain which makes it impossible to falsify the transaction history unless the Bitcoin blockchain can be compromised as well.

The full prototype implementation of our platform and the data used for the experiments is publicly available on GitHub for reproducibility purposes^1^. We note that the open-source components that we used in our implementation have been chosen for their conveniency and proven reliability. However, we emphasize that our approach is tool-agnostic and that each component can be replaced by better and more efficient implementations if available.

#### 2.6 Survey

The survey of citizen concerns in regards to genetic testing and genetic data sharing (Figure 1) was conducted by sending an email to 1991 individuals on a mailing list of Nebula Genomics. The recipients were requested to share their views on genomics data privacy by answering two questions (Figure 1A and 1B). 442 individuals participated in the survey.

### 3 Additional Performance Results

In this section, we describe the experimental setup used for the performance analysis of our platform prototype implementation and we provide additional performance results that are not part of the main text due to space constraints.

#### 3.1 Experimental Setup

The initial testing environment comprises three computing nodes interconnected by 10 Gbps links, featuring two servers types A and B:

- Type A server: Two Intel Xeon E5-2680 v3 CPUs @2.5 GHz that support 24 threads on 12 cores, and 256 GB RAM.
- Type B server: Two Core Intel Xeon Platinum CPUs @3.4GHz and 3.75 GB RAM with the blockchain database stored on a connected Elastic Block Store (EBS) drive.

We used servers of type A for running the core back-end components the secure privacy-preserving component and for storing the encrypted genetic variants and clinical attributes in a PostgreSQL database. We used servers of type B for storing and managing the blockchain.

In order to test the scalability and parallelization of our system, we increased the number of computing nodes from 3 to 12. To set up our system and facilitate its deployment, we use Docker (*60*).

#### 3.2 Dataset

To evaluate the performance of our system we used open access genomic and clinical data from The Cancer Genome Atlas (TCGA) (*61*). The initial dataset contained 1 million genetic variants distributed across 8,000 individuals. We synthetically augmented this initial dataset in order to obtain the final test dataset of 28 billions genetics variants distributed across 150,000 individuals (each individual has a number of genetic variants that ranges from 15,000 to 200,000).

#### 3.3 Performance Evaluation

In addition to the performance evaluation reported in the main text, we run the following additional experiments with each measurement averaged over 10 independent runs:

- *Experiment 1 – Data preparation and loading time per data provider:* Figure S7, shows the time needed for a data provider (individual) to encrypt an increasing number of clinical attributes and genetic variants and measured in a laptop with a Core i7 processor at 2.8 GHz and 16 GB of RAM.

**Figure S7:**
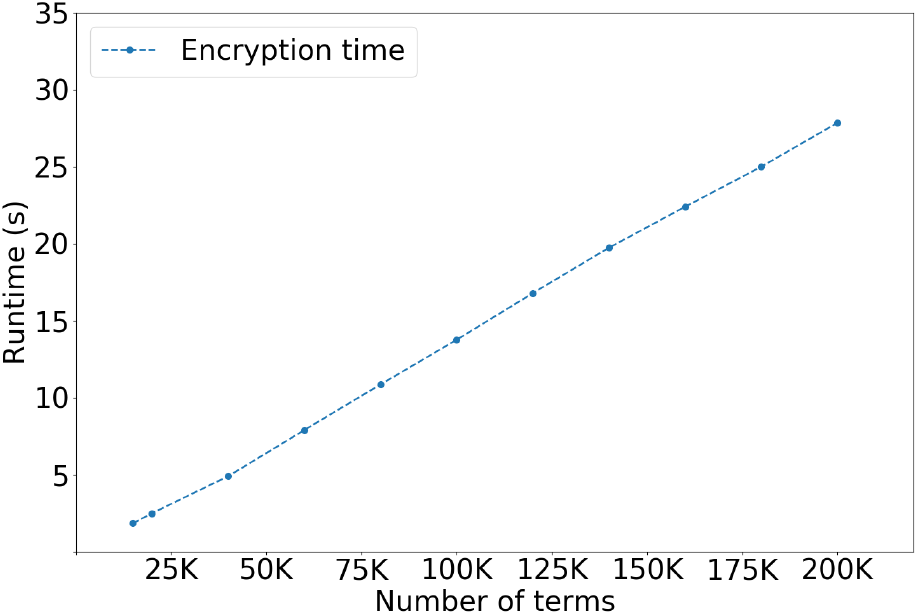
Run time needed for encrypting an increasing number of clinical attributes and genetic variants at a data provider computer (before upload to the system).

Figure S8 shows that the time needed by three computing nodes to re-encrypt the uploaded clinical attributes and genetic variants codes from homomorphic encryption to equality-preserving encryption is linear in the number of codes.

Similarly, Figure S9 shows the time needed to re-encrypt 200,000 uploaded clinical attributes and genetic variants codes from homomorphic encryption to equality-preserving re-encryption is also linear in the number of computing nodes involved in the protocol.

Finally, it is worth noting that the data upload only happens once per data provider and that the overhead introduced by the encryption and the secure re-encryption protocol is always (3 to 4 times) lower than the time it takes to do an insert operation of 1M variables in the database. This time indeed oscillates between 15 and 20 minutes regardless of the data being encrypted or not.

**Figure S8:**
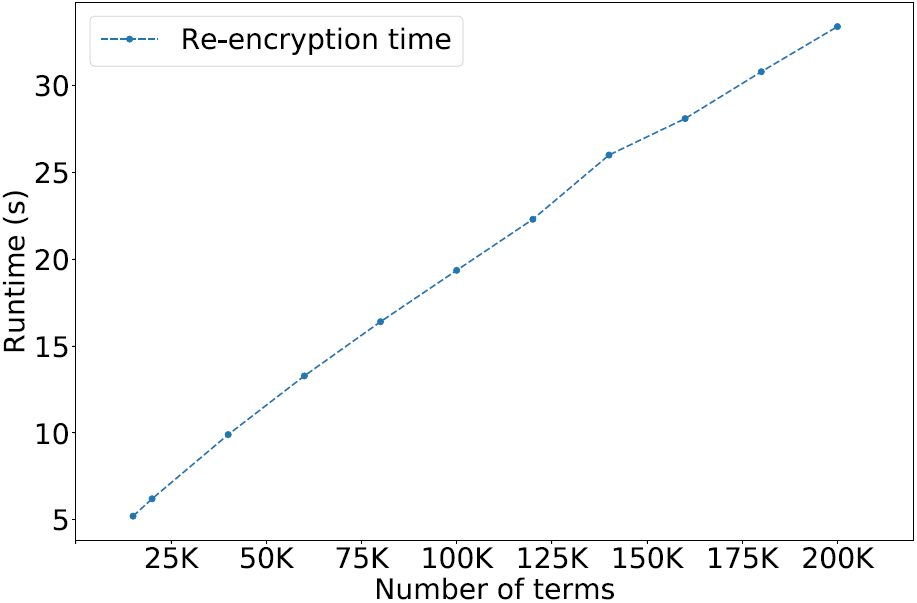
System response time for the upload of encrypted observations from a data provider for a growing number of variables, with three computing nodes.

**Figure S9:**
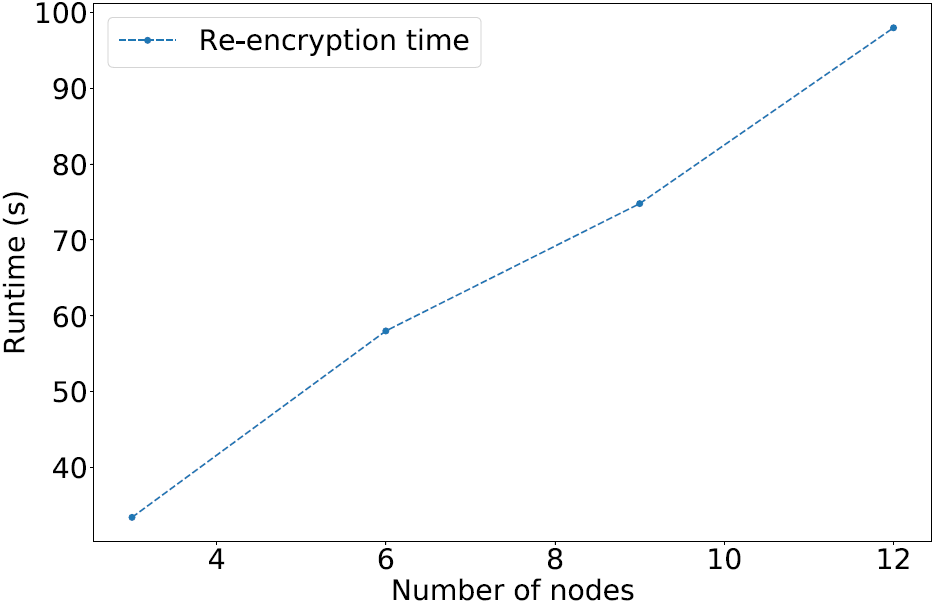
System response time for the upload of encrypted observations from a data provider for a growing number of nodes, with a fixes set of 200k variables (individual variants).

- *Experiment 2 – Query run time breakdown:* Figure S10 explains the almost negligible overhead introduced by the proposed encryption techniques (also shown in Figure 4 of the main text). In particular, we observe that most of the query processing time is consumed by the execution of the query at the database level within the storage unit. This operation is not affected by the encryption, as it takes into account only the time needed to perform a set of standard SQL queries. This time mainly depends on the number of stored observations in the database and of the number of items composing the query.

**Figure S10:**
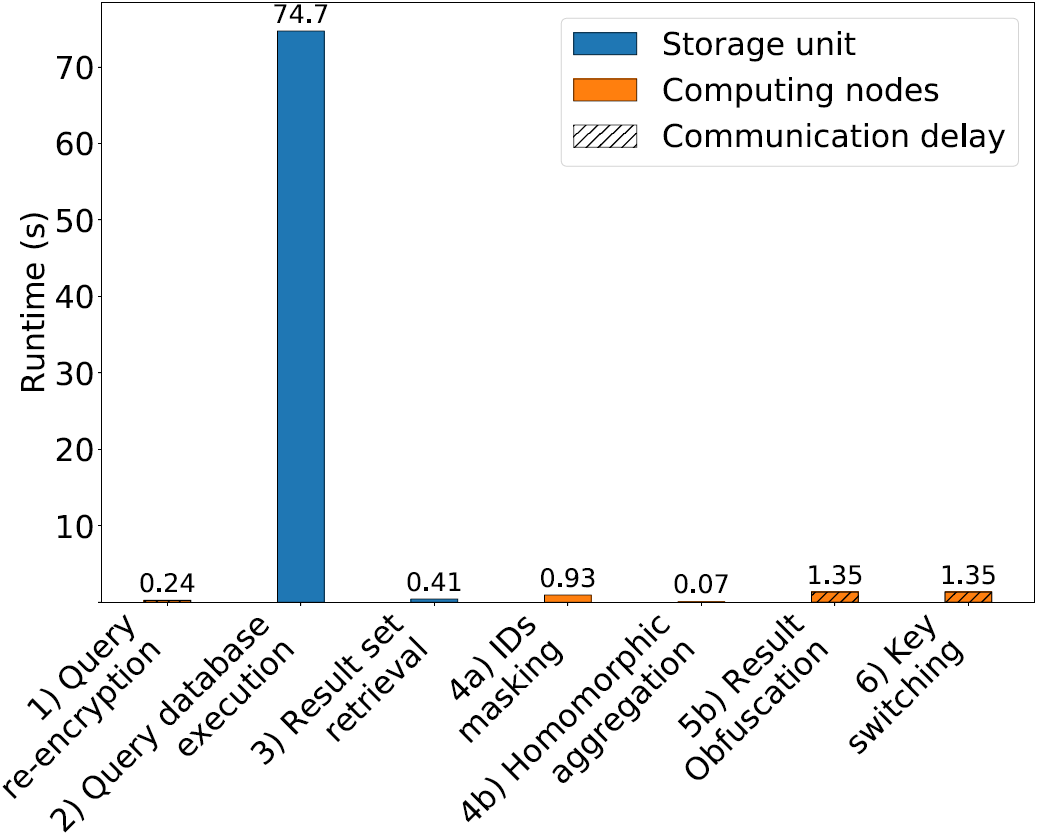
Breakdown of query response time for 28 billion variables, 12 computing nodes and storage units, 10 query variables and with a result set of 1511 individuals per node.

- *Experiment 3 – Blockchain performance:* Figure S11 reports the performance of our blockchain implementation, where the number of transactions per second is only slightly affected by the increase of the number of computing nodes. This demonstrates that the time to generate transactions for data and access policy uploads, for data discovery queries and data access requests is negligible with respect to the time needed to execute these operations.

Finally, the storage overhead introduced by encryption affects only the set of unique observation codes used in the system, and it is in the order of 4x, as the use of equality-preserving converts each observation code, represented by a 64-bit integer, into a 32-bytes ciphertext. Depending on the specific distribution of codes across data providers, a varying number of dummy individuals is also added and must be considered as additional storage overhead. In the tested dataset we obtain an increase factor of 3.6x, which means that for each real data provider, 3.6 dummy individuals (on average) have to be generated.

**Figure S11:**
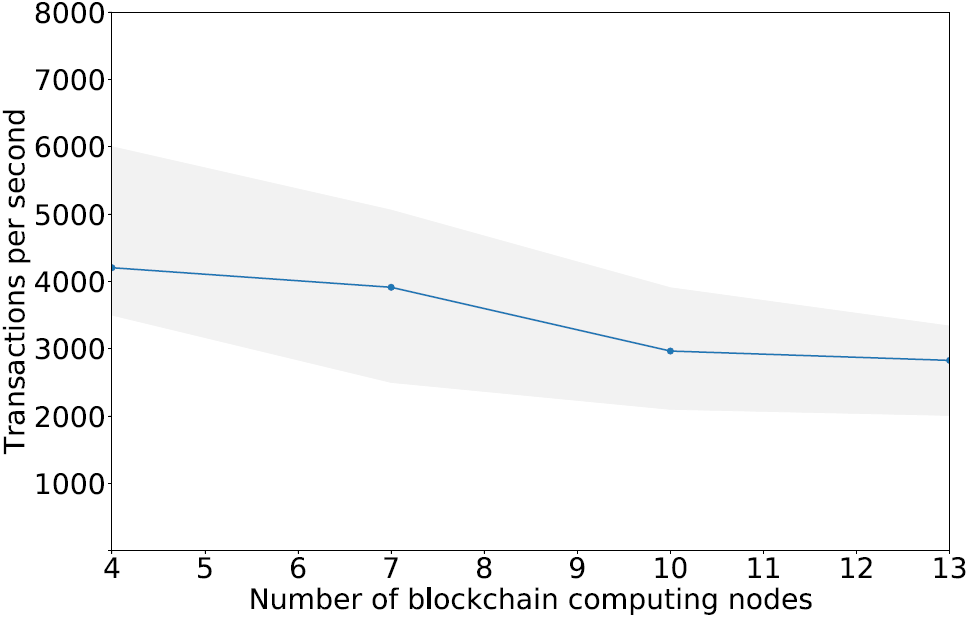
Blockchain performance in terms of number of transactions per second. The shaded area represents lower and upper throughput bounds.

### 4 Security Analysis

The main security and privacy guarantees provided by our platform are described in the main text. These are trust distribution, end-to-end data protection and differential privacy. Here, we briefly discuss and analyze the fulfillment of these guarantees, and we revisit possible extensions for more stringent adversarial models.

The security of our platform is based on the cryptographic guarantees provided by the underlying decentralized protocols described in Section 2.3. All input sensitive variables are encrypted either with equality-preserving encryption (clinical and genetic variable codes) or with homomorphic encryption (individuals’ binary flags) under collectively maintained keys, such that they cannot be decrypted without the cooperation of all sites, thus guaranteeing confidentiality and avoiding single points of failure. For the full step-by-step security analysis of the distributed sub-protocols, we refer the reader to our prior work on the UnLynx library for distributed secure protocols (*33*). Following this analysis, paired with the dummy data strategy described in Section 2.4, it can be seen that our platform protects the query result confidentiality, as only the authorized data queriers can decrypt the query results thanks to the distributed key-switching protocol. Conversely, to avoid re-identification (or attribute disclosure) attacks, our platform also enables the application of differentially private noise to the results and, due to the proposed dummy strategy, it guarantees confidentiality of the data also against the storage unit and all the participating computing nodes.

Our threat model assumes honest-but-curious storage unit and computing nodes, based on the damage to reputation that any of these entities would suffer if it misbehaves in a collective data-sharing protocol. Nevertheless, it is also possible to cope with malicious entities by using protocols (*33*) that produce and publish on the blockchain zero-knowledge proofs for all the computations performed by the computing nodes; the proofs can thus be verified by any entity in order to assess that no party deviated from the correct behavior. This solution yields a hardened and resilient query protocol, but the cost of producing all proofs results in higher and still unpractical computation cost.

## Footnotes

1 https://github.com/ldsec/ccgd-platform

